# Mechanistic Insights into the Substrate Egress Pathways of Human Glucose Transporters GLUT1 and GLUT9

**DOI:** 10.1101/2025.03.04.641472

**Authors:** Manming Xu, Jiwen Jiang, Lin Gao, Saleh O Alyemni, Shozeb Haider

**Affiliations:** UCL School of Pharmacy, University College London, London WC1N 1AX, U.K; University of Tabuk (PFSCBR), Tabuk, Saudi Arabia; UCL Center for Advanced Research Computing, University College London, WC1H 9RL, U.K

**Keywords:** GLUT1, GLUT9, pathways, glucose, urate, apigenin

## Abstract

Glucose transporters (GLUTs) play critical roles in cellular energy homeostasis and substrate-specific transport. Dysfunctional mutations can cause GLUT1 deficiency syndrome and excessive expression of GLUT1 is linked to cancer progression; while abnormal regulation of urate transport by GLUT9 is associated with hyperuricemia and gout. In this study, machine-learning driven molecular dynamics simulations have been employed to investigate the mechanistic insights into the substrate egress pathways of GLUT1 and GLUT9, including the inhibition mechanism of GLUT9 by apigenin. Our findings reveal that intracellular helices play a crucial role in facilitating the transition from inward-closed to inward-open conformations in both transporters. Additionally, aromatic residues, F_291_ and W_388_ in GLUT1, and W_336_ and F_435_ in GLUT9, are identified as key mediators of conformational changes. Analysis of substrate exit pathways provides mechanistic insights into transport profiles and aligns with clinically observed mutations. Furthermore, the inhibitory effect of apigenin on GLUT9 is shown to arise from steric hindrance due to increased substrate size, rather than stable interactions. These findings enhance our understanding of GLUT transporter dynamics and highlight the potential of targeting substrate pathways for therapeutic intervention.

## Introduction

Glucose transporters (GLUTs) are a family of membrane proteins that facilitate the movement of glucose and other substrates across cell membranes, playing essential roles in cellular metabolism and homeostasis^1^. Among the 14 members of the GLUT family encoded by the *SLC2A*, GLUT1 and GLUT9 have attracted significant attention due to their distinct substrate specificities and physiological functions^2^. GLUT1, widely expressed across various tissues, is primarily responsible for basal glucose uptake, which is crucial for energy supply in metabolically demanding organs such as the brain and erythrocytes^2^. In contrast, GLUT9 exhibits unique functional characteristics, extending beyond glucose transport to play a critical role in urate handling. Dysregulation of urate transport by GLUT9 is implicated in hyperuricemia and gout^3, 4^. This has led to a focus on research and development of drugs that target GLUT9 and competitively inhibit its binding to urate, such as Apigenin^5^.

Both GLUT1 and GLUT9 consist of 12 transmembrane α-helices. Interestingly, although GLUT1 and GLUT9 share a significant similarity in structural conformations, they only share a 39.37% sequence similarity (Figure 1). The difference is that GLUT9 is enriched in polar residues and has 5 intracellular helices, whereas GLUT1 is enriched in hydrophobic residues and has only 4 intracellular helices^5–7^. Moreover, compared to GLUT1, which exclusively transports glucose, GLUT9 can transport both glucose and urate but has a 50 times preference for urate^8^. The recent publication of Cryo-EM structures of human GLUT9 in complex with urate and the flavonoid inhibitor apigenin^5^ has elucidated the structural basis and the critical residues involved in urate transportation and apigenin inhibition, highlighting key aspects of ligand binding. However, Shen et al^5^, failed to provide information on the dynamic processes essential for the transporter function, thus limiting our understanding of the specific mechanisms of ligand selection.

**Figure 1.**
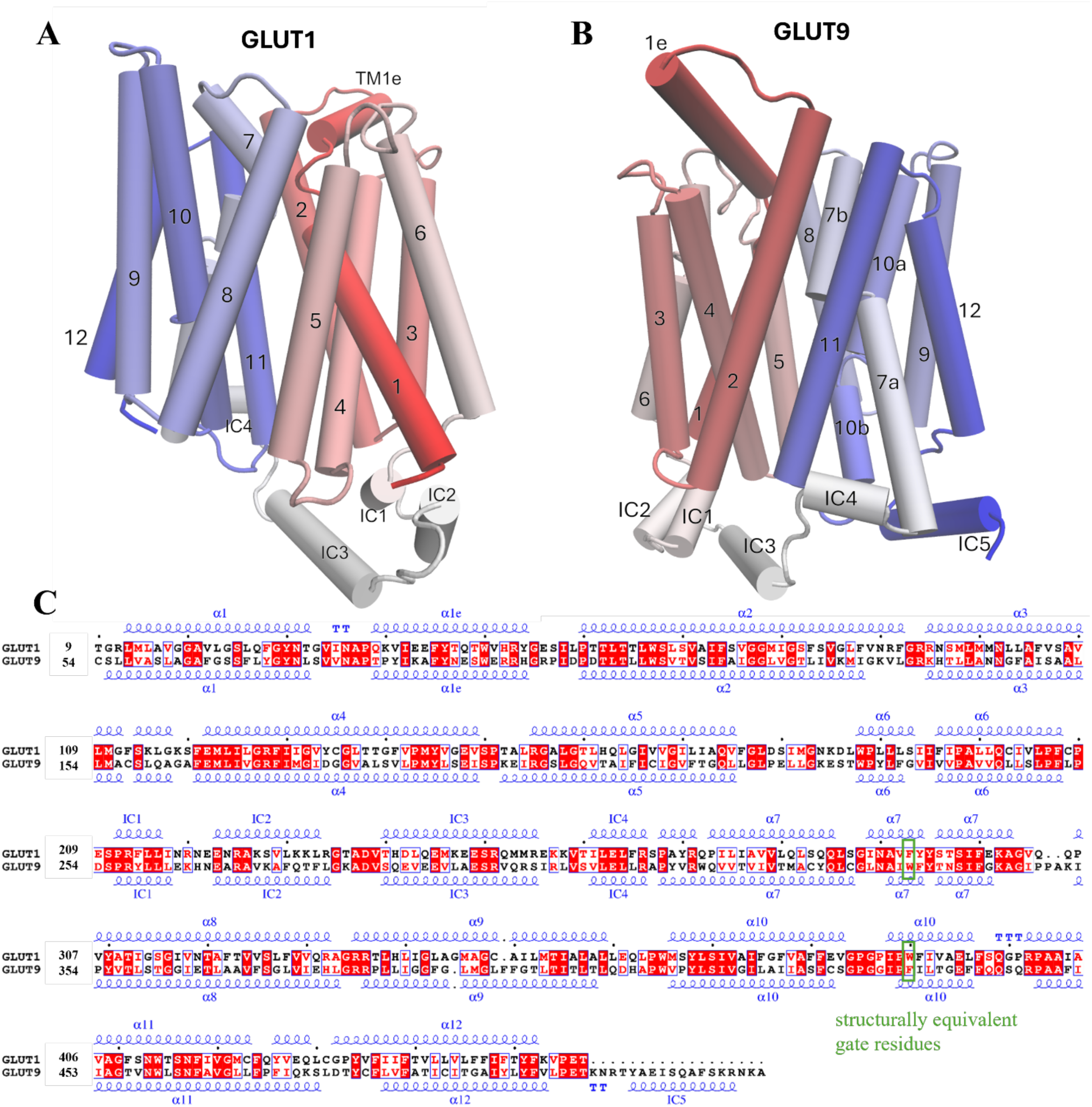
**A.** Structure nomenclature of GLUT1, containing 12 transmembrane helices and 4 intracellular helices. B. Structure nomenclature of GLUT9, consisted of 12 transmembrane helices and 5 intracellular helices. C. Sequence alignment of GLUT1 and GLUT9, with a sequence similarity of 39.37%.

Molecular Dynamics (MD) simulations offer a powerful means to study protein dynamics and protein-ligand interactions, revealing critical conformational changes and key pathways in substrate translocation^9, 10^. Markov state model (MSM) is a method widely used in combination with MD simulations to study the long-time scale conformational dynamics of biomolecules^11^. In some systems, traditional MSMs require long lag times to achieve Markovianity between states. This limits temporal resolution^12^, especially for systems like GLUTs where slow transitions and long-lived intermediate states exhibit non-Markovian behavior. To overcome this challenge, we applied a non-Markovian modeling approach using the recently introduced Integrative Generalized Master Equation (IGME) to investigate GLUT1 and GLUT9 dynamics and apigenin inhibition of GLUT9. IGME incorporates memory effects to accurately model non-Markovian dynamics without the need for long lag times^13^. In conjunction with the IGME, we employed PathDetect**-** SOM, a neural network-based tool that employs Self-Organizing Maps (SOMs) to cluster conformations from MD simulations and map ligand-binding pathways in a 2D space^14^. PathDetect-SOMs allowed us to trace and compare the pathways of urate and glucose in both GLUT1 and GLUT9, identifying key conformational states.

By integrating PathDetect-SOM with IGME, we have achieved a detailed understanding of dynamics and substrate specificity in GLUT9 and GLUT1 across various ligand-bound and inhibitor-bound states. These insights identify molecular determinants driving GLUT9 function, advancing our knowledge of substrate recognition mechanisms within the GLUT family and supporting the development of targeted therapies for metabolic disorders linked to urate and glucose transport dysregulation.

## Results

### non-Markov State Models (nMSM)

Three converged non-Markov state models were constructed for GLUT1 apo, GLUT9 apo, and GLUT9 API (apigenin binding) separately. The feature applied to each system is identical, using the χ_1_ angle of selected binding essential residues and the distance between helices (details in Supplementary). 10 metastable states were extracted for GLUT1 apo at a lag time of 5ns (Figure S1.A and S1.C). State 1 is the highest energy state in GLUT1 and state 10 is the lowest energy state accounts 28.3% of the overall trajectory (Figure S1.D). The flux analysis shows that the most frequent pathway from state 1 to state 10 is: 1-3-7-8-10 (Figure S1.C). The model shows a good alignment with the simulation data in the Chapman–Kolmogorov (CK) test (Figure S1.E), indicating that the nMSM accurately captures the long-timescale dynamics of the system. The mean first passage time (MFPT) was calculated based on the prediction of IGME. MFPT is not fully consistent with the flux network as we are dealing with the non-Markovian process (Figure S1.F).

Similarly, 7 metastable states were separated from the GLUT9 apo system at a lag time of 2ns (Figure S2.A and S2.C), with state 1 being the highest energy state and state 7 being the lowest (Figure S2.D). The flux analysis indicates that the most frequent pathway from state 1 to state 7 passes through state 4, which is a high-energy intermediate (Figure S2.C). The flux result aligns with the estimated MFPT among states (Figure S2.F). For the apigenin binding system of GLUT9, 6 metastable states were observed, with the most frequent pathway detected as 1-4-6 (Figure S3.C) at a lag time of 2ns (Figure S3.A). The MFPT results are consistent with the flux analysis (Figure S3.F). Both models have passed the CK test, confirming their reliability (Figure S2.E and S3.E). A more detailed description of the parameters used to build the nMSM is presented in Methods section.

### GLUT1 Apo Dynamics Highlight the Conformational Role of the ICs, F291 and W388

Metastable states in GLUT1 Apo can be clustered as outwards open, intermediate, and inwards open conformations based on the solvent accessible surface area (SASA) value of the extracellular and intracellular side. The outwards open conformation would have a higher SASA value on the extracellular side and a lower SASA value on the intracellular side and vice versa. The SASA result suggests that the most frequent pathway in this system describes the protein dynamics change from outwards open (state 1) to a relatively stable intermediate (state 8) and eventually reaches the lowest energy inwards open (state 10). State 1 has the largest extracellular side SASA, suggesting that it is more outwards open (Figure 2.A). Conversely, state 10 has the largest intracellular side SASA, implying that it is more inwards open (Figure 2.B).

**Figure 2.**
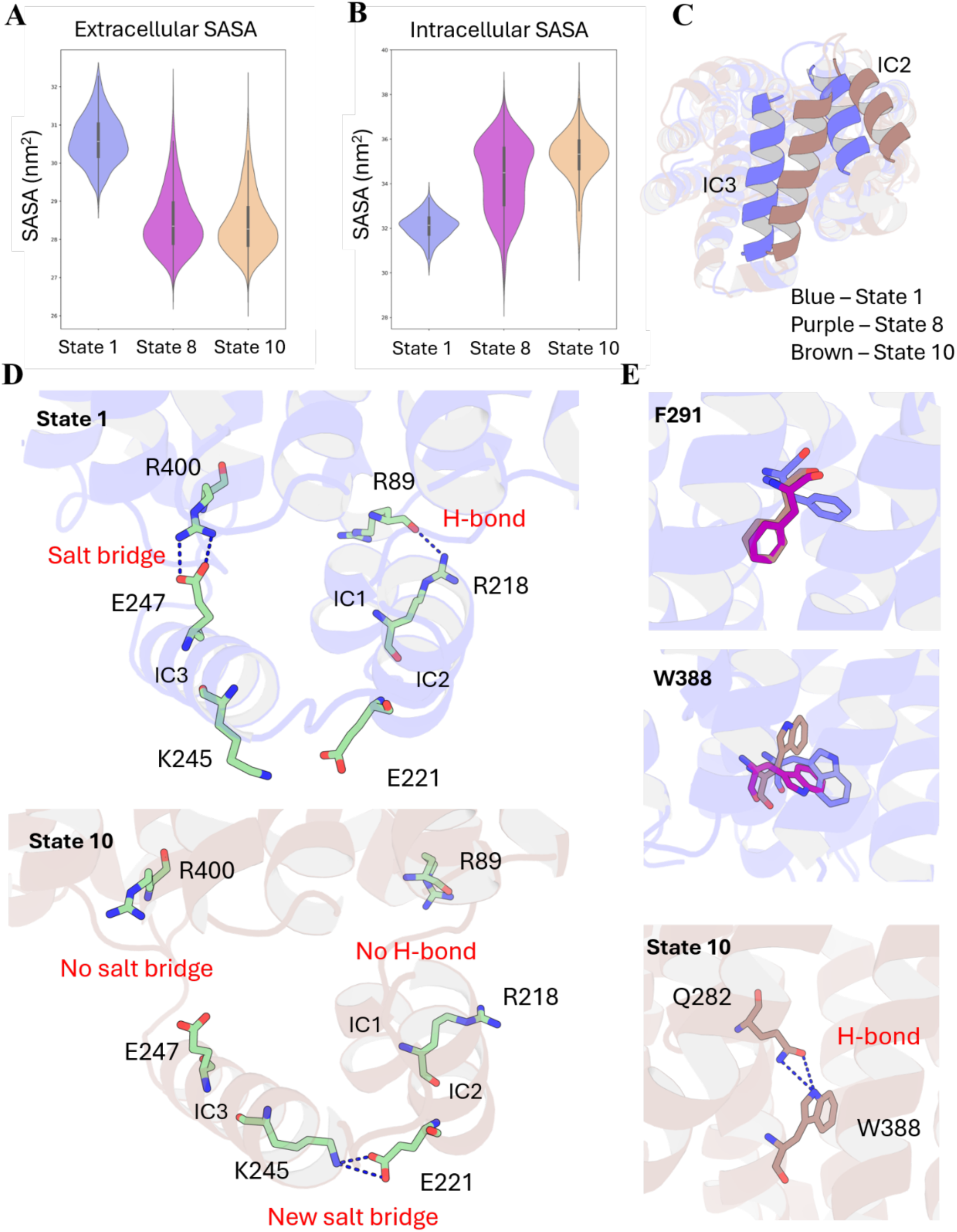
GLUT1 MSM Analysis. **A.** SASA value of extracellular side. **B.** SASA value of intracellular side. **C.** Structure alignment of state 1 and state 10 (bottom view), where IC3 and IC2 undergo significant movements. **D.** In the inwards close state 1, R_400_ and E_247_ can form a stable salt bridge, R_89_ and R_218_ are hydrogen bonds to each other. No interaction can be observed between E_221_ and K_245_. In the inward open state 10. Previously witnessed interactions are completely lost, while a steady new salt bridge is formed between E_221_ and K_245_. **E.** Conformation change of F_291_ and W_388_ among the three states, and the H-bond interaction formed between Q_282_ and W_388_ in state 10. Blue – State 1, Purple – State 8, Brown – State 10.

Superimposition of state 1 and state 10 illustrates significant movements among the intracellular helices (ICs), especially IC2 and IC3 (Figure 2.C). In state 1, IC3 blocks the tunnel, while in state 10, IC3 flips remarkably and leads to the opening of the intracellular side (Figure 2.C). Thus, understanding the movement of these ICs becomes essential as they correspond to the open and close dynamics. Several reasons have been identified triggering the motion of IC3. Firstly, the loss of interaction between IC3 and TM11 frees IC3 and enables its further movements. R_400_ (TM11) and E_247_ (IC3) form a strong salt bridge and hydrogen bond interactions in state 1 (Figure 4.D). Distance calculation indicates that the sidechain between these two residues are within 4Å in state 1 over 80% simulation time (Figure S4.A and S4.B). However, these interactions are completely lost in state 10 (Figure 4.D), where the residence time of this interactions decreases to approximately 5% (Figure S4.A and S4.B). Moreover, the flip of IC1 makes more space and contributes to the movement of IC3. IC1 maintains its position in state 1 through a hydrogen bond interaction between the R_89_ backbone (TM2) and the R_218_ sidechain (IC1) (Figure S4.D) with over 60% residence time (Figure S4.C). In state 10, this hydrogen bond is much less likely to occur (Figure 4.D and Figure S4.C). Lastly, the strong salt bridge and hydrogen bond between K_245_ (IC3) and E_221_ (IC2) drags IC3 to move towards the N terminal of helix 1, leading to the significant movement of this secondary structure (Figure 4.D). The carboxylate group of E_221_ show very close contact with sidechain χ-nitrogen of K_245_ in state 8 and state 10, based on the distance calculation, indicating strong interactions (Figure S4.D and S4.E). These factors collectively drive IC3 to move closer to the N terminal of helix 1, leading to the opening of the intracellular side (Figure 4.D).

Furthermore, two residues with aromatic ring structures are identified as essential for the transporter^15, 16^. F_291_ is situated at the extracellular gate. The χ_1_ angle distribution of F_291_ indicates that the two high energy states (state 1 and state 2) prefer a trans(180°) orientation, which is significantly different from other states (Figure S5.A). All the other 8 states, which are more energetically favourable, have a gauche(+)(−60°) conformation (Figure S5.A). This results in a different orientation of F_291_ (Figure 2.E). F_291_ has been identified in previous studies as a tracker of glucose^15^, as mutations on this site significantly alter the glucose transport ability of GLUT1^17^. The horizontally placed sidechain in state 1 enables the tracking of glucose via the aromatic ring, indicating that the extracellular side is actually open. While in state 8 and state 10, F_291_ sidechain is placed alongside the TMs (resting state), losing the ability to attract glucose from the extracellular environment. On the intracellular side, W_388_ has been found to be essential for the conformation change from state 8 to state 10, whose χ_1_ angle corresponds to the separation in the free energy plot (Figure S5.A). W_388_ in previous studies has been shown to be essential for the conformational change of GLUT1 from outwards open to inwards open^16^. In state 1 and state 8, the χ_1_ angle of W_388_ is around trans(180°), while in state 10 it changes to gauche(+)(−60°) where the sidechain of the residue rotates resulting in the opening of the tunnel (Figure 2.E). Q_282_ was found to significantly contribute to this orientation of W_388_ by forming strong hydrogen bonds that stabilises the W_388_ sidechain (Figure 2.E). A remarkably residence time of this interaction in state 10 is confirmed by the distance calculation (Figure S5.B). Moreover, W_388_, together with P_141_, M_142_, and I_404_, form a hydrophobic gate that blocks the tunnel in state 1. The rotation of the W_388_ sidechain sterically pushes the residues apart in state 10, thereby opening the tunnel’s inward side (Figure S5.C).

### GLUT9 Apo Dynamics Reveal Similar Dynamic Behaviours to GLUT1

The partial SASA value is used to define the outwards open and inwards open conformation in GLUT9. The SASA value result indicates the nMSM dominant pathway (1-4-7) describes the transition from an inward close conformation (state 1) to an inward open conformation (state 10), as state 10 has a significantly higher value of intracellular SASA (Figure 3.B). However, unlike GLUT1 which witnesses an inverse in the SASA value of the two sides, the opening and closure of the extracellular side in GLUT9 does not seem to affect its SASA value (Figure 3.A).

**Figure 3.**
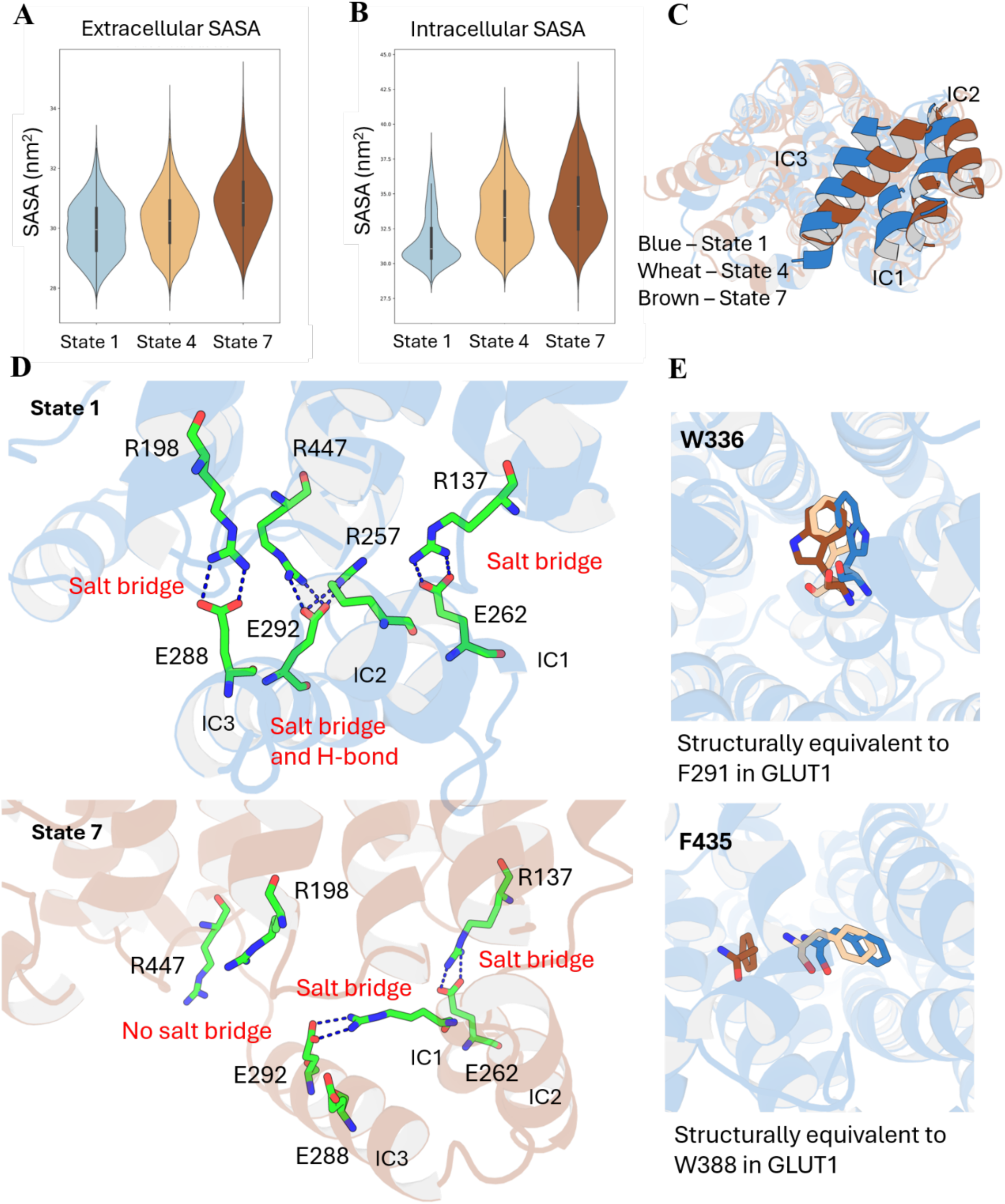
GLUT9 MSM Analysis. **A.** SASA value of extracellular side. **B.** SASA value of intracellular side. **C.** Structure alignment of state 1 and state 7 (bottom view), where ICs undergo significant movements. **D.** In the inward close state 1, four salt bridges could be observed: R_137_-E_262_, R_198_-E_288_, R_257_-E_292_, E_292_-R_447_. In the inward open state 7, salt bridges R_198_-E_288_ and E_292_-R_447_ decrease occurrence, while the other two salt bridges preserve. **E.** Conformation change of W_336_ and F_435_ in each state. Blue – State 1, Wheat – State 4, Brown – State 7.

Similar to GLUT1, GLUT9 undergoes tremendous movement in its ICs, especially IC3, where they move closer to the N terminal of helix 1, and open the intracellular side (Figure 3.C). In state 1, the ICs are held and stabilized through several salt bridges with the protein backbone (Figure 3.D). E_262_ on IC1 forms a stable salt bridge with R_137_ on TM3 (80% residence time) (Figure S6.A). Two salt bridges between E_288_ (IC3) and R_198_ (TM5), and E_292_ (IC3) and R_447_ (TM11) stabilises IC3 in its place in state 1, both with a residence time >60% (Figure S6.B and S6.C). One additional salt bridge between R_257_ and E_292_, links IC1 and IC3 (Figure S6.D). In state 10, however, the residence time of the two salt bridges between the TMs and IC3 drops significantly, which frees IC3 and enables its further motion (Figure S6.B and S6.C). In contrast, the salt bridges that connect IC1 to TM3 and IC3 to IC1 are preserved (Figure S6.A and S6.D). Thus, with the opening of the intracellular side, IC1 would be dragged towards the N terminal of helix 1, which subsequently triggers the movement of IC3 in the same direction, ultimately leading to tunnel opening(Figure 3.D).

Like in GLUT1, we identified two aromatic residues in GLUT9 acting as gate modulators. W_336_ and F_435_, which are structurally equivalent to F_291_ and W_388_ in GLUT1, undergo significant conformational changes along the pathway (Figure 3.E). Unlike F_291_, which is more likely to be a binding residue, W_336_ acts as a gating residue in GLUT9. Urate is much larger than glucose and is prevented from entering the tunnel while the sidechain of W_336_ is positioned perpendicular to the transport axis (Figure S7.A and S7.B). Thus, state 4 and state 7, where the trans(180°) conformation of W_336_ is observed, are considered as outwards closed, while state 1, where gauche(-)(60°) conformation dominates is considered as outwards open (Figure S7.C). A very similar phenomenon could also be observed at the intracellular side of GLUT9, where F_435_ modulates the opening and closure of the intracellular side. In this case, state 1 prefers a χ_1_ angle of trans(180°), where the sidechain blocks the tunnel, while state 7 tends to have a χ_1_ angle of gauche(-)(60°), where the intracellular side opens (Figure S7.D). State 4, however, shows a relatively equal preference for both orientations, indicating it is an intermediate state. The significant difference in the sidechain orientation as well as the position from state 1 to state 7 enlarges the tunnel on the intracellular side (Figure S7.E). The observation of the difference in the conformation of these two aromatic residues aligns with the result of nMSM, indicating the pathway successfully describes the conformation change of GLUT9, as well as emphasizes the significant contribution of these two residues to GLUT9 apo dynamics.

### GLUT1 Substrate Exits Pathway Explains Its Preference to Glucose

For the exit of glucose, seven clusters were identified by PathDetect-SOM^14^. According to the distance heatmap, glucose shows the least contact with the selected residues in cluster A, while the most contact in cluster E (Figure 4.A). Thus, cluster E is considered the initial binding state and cluster A represents the completely unbound conformations. Two possible pathways, all starting from cluster E to cluster A, are illustrated in the neuron transition network: E (Purple)-C (Red)-B (Green)-A (Blue), and E (Purple)-D (Orange)-F (Brown)-G (Sky Blue)-A (Blue) (Figure 4.B). At starting point in cluster E, glucose forms close contact with residue P_26_, Q_161_, Q_283_, N_288_, Y_292_, F_291_, N_411_, and N_415_, which is consistent with the conformation observed in the crystal structure (PDB id: 4PYP) (Figure S8.A).

**Figure 4.**
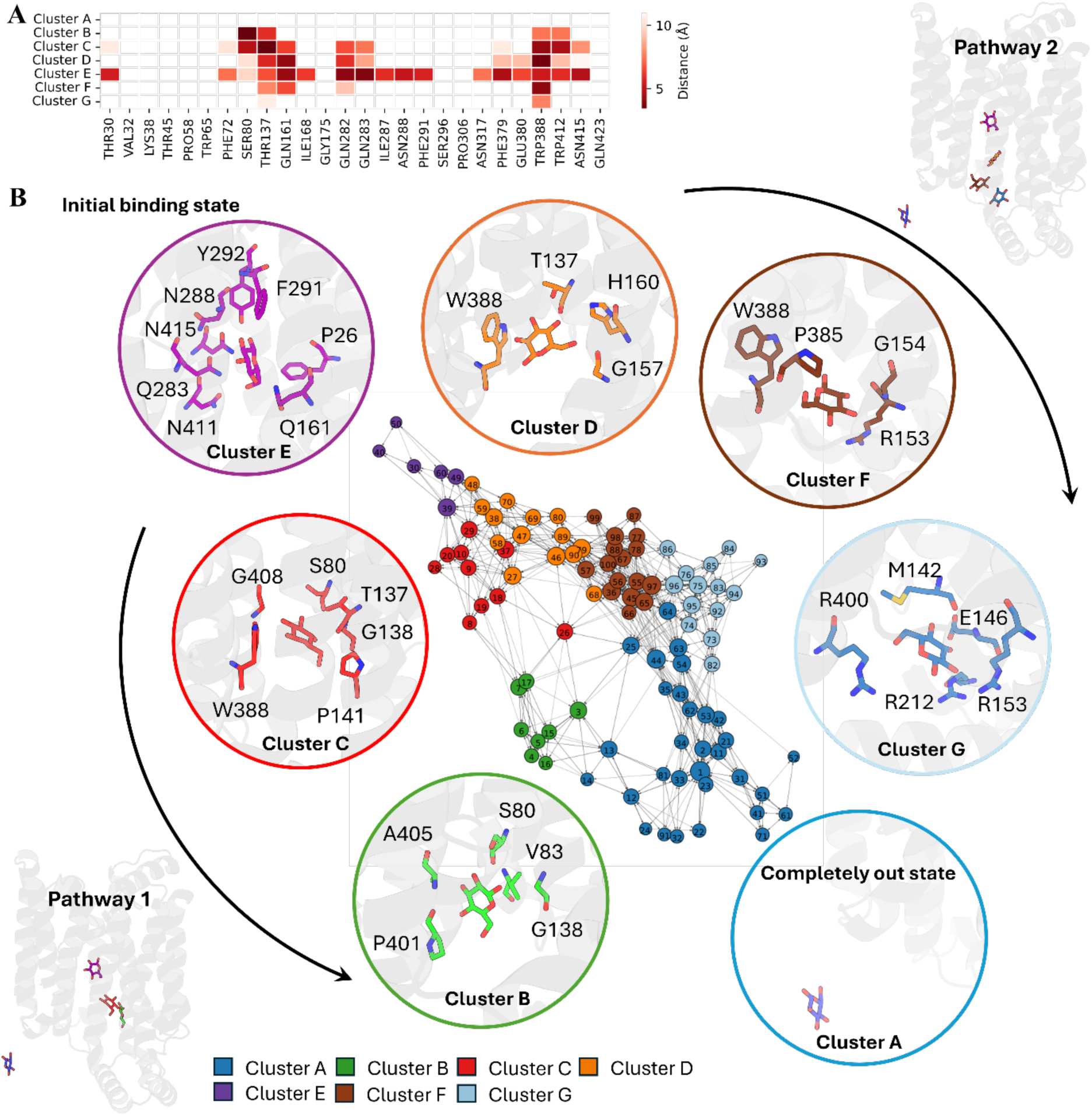
GLUT1 Glucose Pathway Results. **A.** Distance heatmap between glucose and selected input residues during SOM training. **B.** Transition network with potential pathways indicated by black arrows. Detailed ligand-protein contacts in each cluster are placed closed to the neuron with residues within 4Å to the substrate highlighted, each pathway follows the arrow from cluster E to cluster A.

In the first pathway, glucose moves out from the crevice between TM2 and TM11 (Figure S8.B). In cluster C, S_80_, T_137_, G_138_, and W_388_ form contacts with glucose (Figure 4.B). After passing S_80_, glucose enters a site surrounded by residues with short sidechains: V_83_, G_138_, and A_405_ (Figure 4.B). Thus, the protein is not able to hold glucose, and the glucose exits in the next stage. This exit position does not seem to be influenced by the movements of the ICs but depends on the distance between TM2 and TM11 at the intracellular side. In the inward closed structure defined by nMSM previously (State 1), the crevice between TM2 and TM11 is quite narrow, thus eliminating the possibility of the escape of glucose (Figure S8.C). An alternative exit position is also observed in pathway 2, where glucose moves out of the tunnel opened by the movement of IC3 (Figure S8.D). Glucose first shows very close contacts with T_137_, G_157_, H_160_, and W_388_. After cluster D, glucose slides vertically to a hydrophobic environment formed by R_153_, G_154_, P_385_, and W_388_. It then follows a relatively polar path surrounded by M_142_, G_145_, E_146_, R_153_, R_212_, and R_400_. Unlike the first pathway, this route is strongly modulated by the movement of the ICs, especially IC3. Superimposition of the structures indicates that glucose is not able to exit the protein through this pathway in the inward closed conformation defined by the nMSM model (state 1), as the tunnel is blocked by IC3 (Figure S8.C). W_388_, an important residue we mentioned in GLUT1 apo dynamics, contributes to both glucose pathways with its sidechain flipped up, opening the intracellular gate as we proposed (Figure 4.B).

Compared to glucose, urate is much larger and contains one additional ring structure. In nature, the GLUT1 transporter does not transport urate^5^. Ten clusters were identified when analysing the urate exit pathway of GLUT1 (Figure S9). From the distance heatmap, cluster H is identified as the unbound state (Figure S9.A). However, to identify the bound state of urate, where in clusters C, D, and E all show close contacts between the substrate and the protein, was challenging. From the neuron transition network, cluster D (Orange), which is located at one edge of the network, is assumed to be the initial binding stage (Figure S9.B). Unlike glucose, the transit of urate does not follow a clear route to exit as it tumbles inside the tunnel (Figure S9.B). Cluster B and Cluster F are indistinguishable and display an unstructured transition network (Figure S9.B). Detailed contact conditions in these clusters suggest that the urate is held in a very similar place in these two clusters, where W_388_ can form a strong π-π interaction with the urate (Figure S9.B). This interaction is not observed in the glucose system, as glucose lacks an aromatic ring and does not have the delocalized π-electron system required for the interaction. Thus, the rotation of the sidechain of W_388_ only opens the tunnel in the glucose system and does not strongly influence its exit. Compared to the glucose system (glucose cluster A), the number of frames in which urate is in the completely unbound state (urate cluster H) is more than three times lower, suggesting a much slower exit. The unstructured pathway of urate in GLUT1 explains why GLUT1 is not able to transport urate.

### GLUT9 Ligand Exit Pathway Explains Its Preference to Urate

Unlike GLUT1 which exclusively transports glucose, GLUT9 can transport both glucose and urate but has a 50 times preference for urate^8^. Thus, understanding why this preference happens is quite essential in understanding the transport mechanism of GLUT9. Pathway analysis provided a clear exit pathway for urate. Detailed contact information illustrates constant interactions between urate and GLUT9 (Figure 5). PathDetect-SOM deduced two unique binding conformations for urate in GLUT9, cluster B and cluster D. In cluster B, the sidechain of R_171_ interacts with urate via hydrogen bonding. Residues with aromatic ring structures appear in the neighbourhood but do not participate in any significant interactions. Another binding conformation, which is more favourable, is cluster D, which shows more stable interactions. W_336_, an essential residue acting as a gate on the extracellular side of GLUT9, forms a π-π interaction with the indole ring of the urate. Moreover, N_79_ and E_364_ forms hydrogen bonds with urate. GLUT9 favours cluster D, as the frames belonging to cluster D are three times more than that of cluster B. Urate then follows a single exit pathway: A (Blue)-C (Red)-E (Purple)-F (Brown)-G (Sky Blue). After passing W_336_, the urate forms hydrogen bond interactions with the sidechain of Y_327_, Q_328_, N_333_, W_336_, and E_364_ (Cluster A). In the next stage (Cluster C), all the hydrogen bond interactions are lost except for that with Y_327_. F_435_ then takes part in and forms a π-π interaction to urate. F_435_ is structurally equivalent to W_388_ in GLUT1. However, the interaction it forms might be weaker compared to tryptophan as it lacks the same electronic complexity as the indole ring of tryptophan. Such a relatively weaker response might allow GLUT9 to balance the role of directing urate and transporting it into the cell. After passing F_435_, the urate then loses all the interactions with the protein in cluster F, thus going out of the transporter in the next stage.

**Figure 5.**
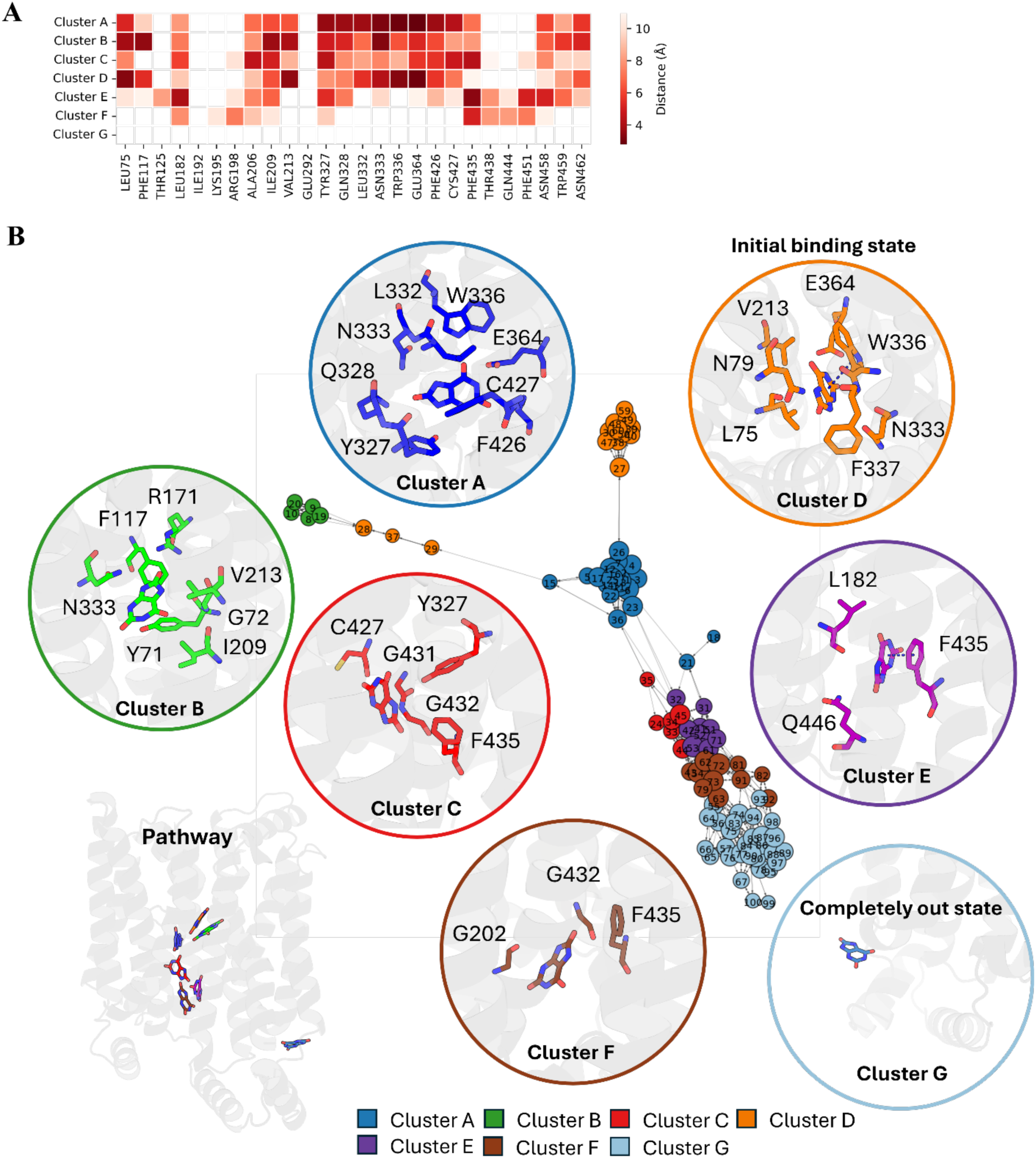
GLUT9 Urate Pathway Results. **A.** Distance heatmap between glucose and selected input residues during SOM training. **B.** Transition network. Detailed ligand-protein contacts in each cluster are placed close to the neuron with residues within 4Å of the substrate highlighted, pathway starts from cluster D and ends at cluster G.

Compared to urate, glucose is smaller and more hydrophilic, surrounded by oxygens that easily form interactions with the polar sidechains alongside the tunnel. As mentioned previously, GLUT9 is more enriched in hydrophilic residues compared with GLUT1. The increased potential interactions between GLUT9 and glucose due to increased substrate hydrophilicity might be one of the reasons to explain the preference of GLUT9 to urate. Nine clusters were defined by PathDetect-SOM, and the distance heatmap indicates a very close contact between glucose and GLUT9 in clusters A, B, C, and D. Cluster I is considered as the completely unbound state for glucose according to the heatmap (Figure S10.A). Acting as another edge of the transition network, cluster B is defined as the starting bound conformation of glucose. Unlike urate which exhibits a clear exit pathway, the transition network of glucose is much more complex, indicating multiple interaction sites inside the tunnel. In cluster B, polar sidechains of Y_71_, Y_327_, Q_328_, and N_333_ are able to form interactions with glucose, which is quite similar to what happens in urate cluster A. However, compared to the urate case, the interactions between glucose with W_336_ or E_364_ are completely lost in the initial state. After leaving the starting state, rather than following a direct pathway, glucose tumbles inside the tunnel. Its position in clusters A, C, and D are quite similar (Figure S10.C). Moreover, in cluster B to cluster A or cluster B to cluster C, glucose moved slightly closer to the extracellular side. Q_328_, N_333_, W_336_, and E_364_, which are essential for the exit of urate, also play a role in the glucose pathway. These residues form interactions with glucose throughout the first several clusters (A, B, C, and D). However, as glucose only contains oxygen, some interactions formed by these residues are much weaker compared to those with urate. This might be one of the plausible reasons that cause the differences in substrate preference. Cluster E becomes a critical intermediate state that connects the four initial clusters (A, B, C, and D) and the four subsequent clusters (F, G, H, and I) (Figure S10.B). Detailed conformation analysis implies a relatively hydrophobic environment within cluster E, with glucose surrounded by hydrophobic residues: L_182_, P_186_, P_434_, F_435_, and I_436_. After passing cluster E, glucose continuously forms close contacts with the protein until it leaves in cluster I. However, urate is much less likely to be sequestered in complex sites, like glucose, after leaving its initial binding state, which would explain the structured exit pathway of urate. Moreover, π-π stacking plays an important role in guiding urate. F_435_, which we proposed as a gate residue having a similar function as W_388_ in GLUT1, forms π-π with urate, providing stability to prevent its rotation inside the tunnel. In contrast, the π-π stacking would all be lost in the case of glucose, resulting in the tumbling and rotation of the substrate around F_435_ as shown in clusters E, F, and H.

### GLUT9 API Dynamics Explains the Inhibition Mechanism of API

Apigenin binding GLUT9 was simulated to study the dominant binding position of this inhibitor inside GLUT9. The most dominant pathway in this system is 1-4-6. Superimposition between state 1 and the Cryo-EM structure shows a very similar apigenin binding position (Figure 6.A), indicating the Cryo-EM binding state is our starting point in this deduced pathway. The most significant differentiation on the free energy map from left to right is caused by the χ_1_ angle of W_336_, an important residue that also undergoes significant conformation changes in GLUT9 apo dynamics (Figure 6.B). The horizontally positioned sidechain of W_336_ in state 1, state 2, and state 4 blocks the tunnel on the extracellular side, indicating an extracellular closed conformation (Figure S11.A). This χ_1_ angle, however, prefers a completely different orientation in the other three states, where the sidechain flips down towards the intracellular side and opens the extracellular side (Figure S11.A). Thus, these states are more likely to be an outward open conformation. The apo dynamics of GLUT9 indicate that the transporter prefers an inwards open conformation, as both the Cryo-EM structure and the extracted lowest energy conformation from nMSM are all inward open. However, with the movement of Apigenin, the transporter transforms from an energetically preferred inward open conformation to a high energy outward open conformation, implying the inhibition of Apigenin.

**Figure 6.**
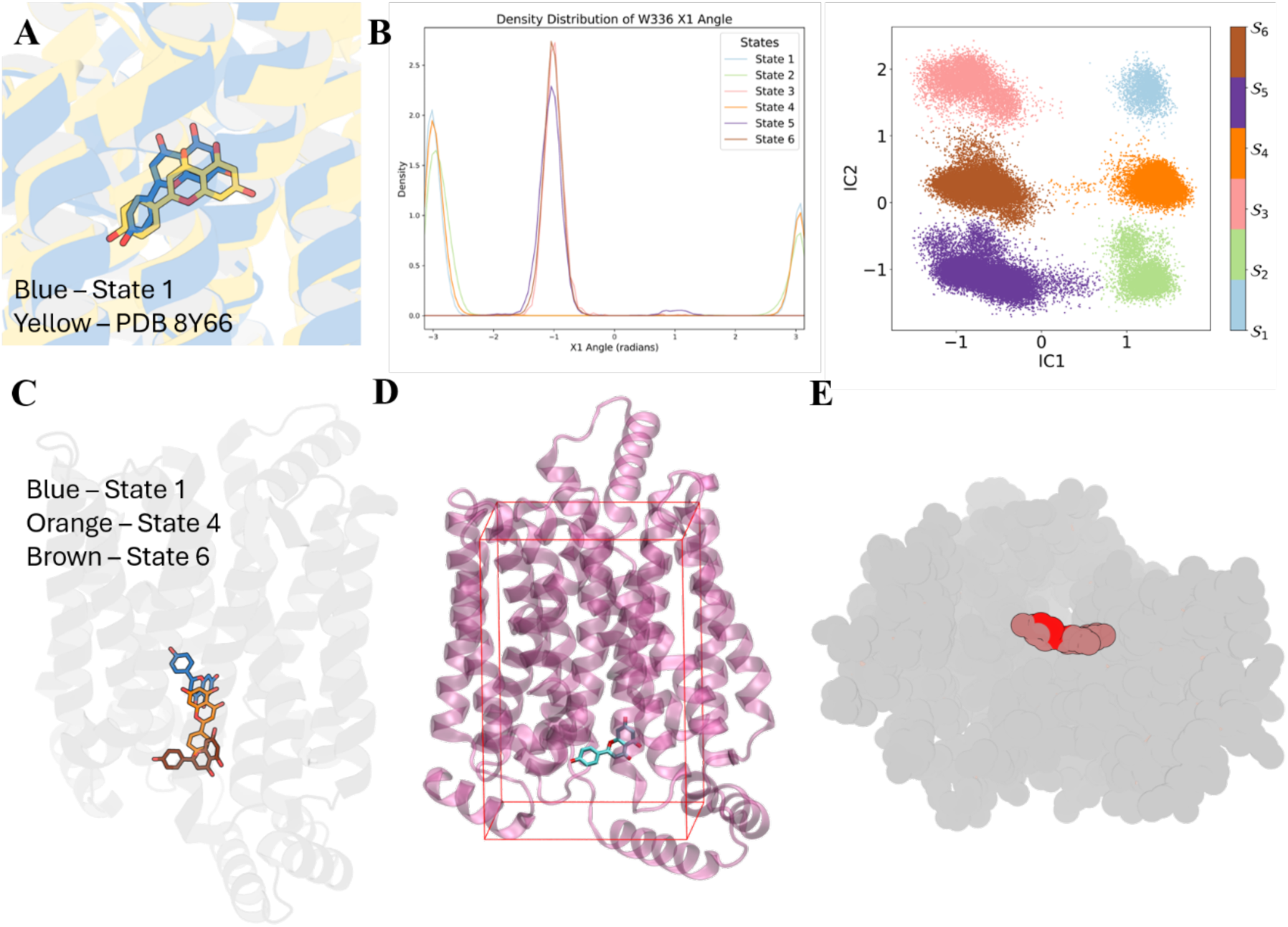
**A.** Superimposition of extracted state 1 and original PDB 8Y66. **B.** The Х_1_ angle of W_336_ and the corresponding state distribution map. **C.** Ligand pathway. **D.** Defined flat-bottomed restraints and extracted structure from state 6. **E**. Spheres representation of GLUT9 and API.

Interestingly, unlike urate and glucose exiting the GLUT9 tunnel during simulations, apigenin was retained within the GLUT9 (Figure 6.D). We initially hypothesized that GLUT9 might form more and stronger interactions with apigenin potentially via multiple hydrogen bonds. To explore this, we analysed the atom-atom distances between GLUT9 and apigenin, seeking possible dipole-dipole attractions. However, no pair of atoms maintained a proximity of less than 4 Å with occupancy above 50%, suggesting that no consistent, close-contact interactions primarily account for apigenin’s retention (Figure S11.B). Therefore, we attribute this retention to the macroscopic structural characteristics of the GLUT9-apigenin complex. We propose that the relatively large size of apigenin acts as a plug and prevents its release from the intracellular side. Spheres representation of the extracted structure of state 6 confirms our proposition where Apigenin is clearly a plug-in for this transporter and is unable to escape due to its increased size (Figure 6.E)

## Discussion

### GLUT1 and GLUT9 Share Similar Dynamic Behaviours

GLUT1 and GLUT9 are structurally and dynamically similar to each other. In both proteins the ICs, especially IC3, determine the opening of the intracellular side through the modulation of a series of salt bridges formed either between ICs or between ICs and TMs. Both proteins contain two dynamically critical aromatic residues, F_291_ and W_388_ in GLUT1, and W_336_ and F_435_ in GLUT9, which are structurally equivalent. F_291_ in GLUT1 tracks glucose and thus needs to orient in a trans(180°) conformation to expose its sidechain to the solvent. On the other hand, W_388_ in GLUT9, W_336_, and F_435_ in GLUT9 act as gate modulators that strongly correspond with the open and closed conformations on each side.

### Pathway Detection Explains Substrate Preference and Links with Clinical Studies

Glucose and urate pathways in each protein have been analysed, revealing distinct substrate preferences. GLUT1 is incapable of transporting urate, whereas GLUT9 demonstrates a 50-fold preference for urate over glucose^8^. The desired substrate in each case presents a structured pathway. Despite the dynamic similarities between the two proteins, differences in the chemical profile of the transport tunnel and the intracellular exit site significantly influences substrate selectivity. In GLUT1, the potential strong π-π interaction formed between W_388_ and urate disrupts the urate pathway. In contrast, the substitution of W_388_ with F_435_ in GLUT9 reduces electronic complexity, weakening the interaction with urate. This alteration facilitates the smoother release of urate while providing additional stabilization that minimizes substrate misorientation. However, the present study focuses primarily on substrate exit pathways, leaving the extracellular entry mechanisms underexplored. As substrate recognition and uptake are equally critical for transport efficiency, future investigations on substrate entry dynamics would be highly valued.

Moreover, clinically detected mutations that impact the transport ability of both transporters were identified alongside the recognized pathways. Missense mutations on T_137_, R_153_, E_146_, and M_142_ have been isolated from patients with GLUT1 deficiency syndromes^18^. These residues are shown to be directly involved in the glucose pathway, as all of them appear in cluster G in the glucose pathway. For GLUT9, substitutions on L_75_, R_171_, and N_333_ have been identified, where mutations on L_75_ and N_333_ harm the urate uptake significantly, while mutation on R_171_ gives a relatively moderate effect^19^. The identified pathways in GLUT1 and GLUT9 offer valuable insights into the mechanistic impacts of these mutations, enhancing our understanding of how specific residues influence substrate transport and their potential roles in disease pathology.

### API Acts as a Plug to GLUT9

Apigenin is a natural product that has been found as a GLUT9 inhibitor^20^. Previous studies and the published Cryo-EM structure indicate several hydrogen bond interactions that anchor apigenin inside the transporter^5^. However, based on our dynamic analysis, no consistent interactions could be formed between GLUT9 and apigenin. The sphere representation of the lowest energy structure suggests that apigenin inhibits the transporter purely due to its size and acts as a plug that blocks the GLUT9 tunnel.

## Conclusions

In conclusion, our study compares the apo dynamics of GLUT1 and GLUT9, emphasizing their dynamical similarities. We analysed the exit pathways for glucose and urate in both transporters, explaining the preference of urate to glucose in GLUT9, as well as the inhibition mechanism of GLUT9 inhibitor apigenin. These findings enhance our understanding of the substrate selectivity and regulatory mechanisms of these transporters, especially for GLUT9. Future studies integrating the extracellular entry pathways could further refine our understanding of substrate recognition, transport dynamics, and inhibition strategies, paving the way for improved therapeutic targeting of GLUT transporters.

## Methods

### Building the GLUT systems

The cryo-EM structures of the membrane proteins GLUT1 (PDB :4PYP) and GLUT9 (PDB: 8Y65) were obtained from the PDB. HTMD python package^21^ was used to build seven systems (details in Table 1) as follows GLUT1-Apo, GLUT1-Glucose (with glucose in the binding site), GLUT1- Urate (with urate in the binding site), GLUT9-Apo, GLUT9-Glucose (with glucose in the binding site), GLUT9-Urate (with urate in the binding site) and finally GLUT9-API (with apigenin in the binding site). The protonation states of the titratable residues were predicted based on pH 7 using PROPKA3 and PDB2PQR as implemented in the HTMD function proteinprepare()^22^. A pure 1- palmitoyl-2-oleoylphosphatidylcholine (POPC) membrane 130 x 130 Å was built using the membrane builder class in HTMD. After that, each of the prepared proteins was embedded into a separate membrane. The systems are then solvated by adding a 20 Å layer of water molecules above and below the membrane and then neutralised to a 0.15M NaCl concentration and built using the charmm. builder in HTMD. The proteins, ions, and lipids were parameterized using the CHARMM36m^23^ forcefield, and TIP3P^24^ was used as the water model. The ligands were parameterized using the CGENFF^25^. Firstly, the correct mol2 file for each ligand was generated using Schrodinger suite (Schrödinger Release), then the mol2 files were uploaded to the CGENFF web server to generate the parameters.

**Table 1.**
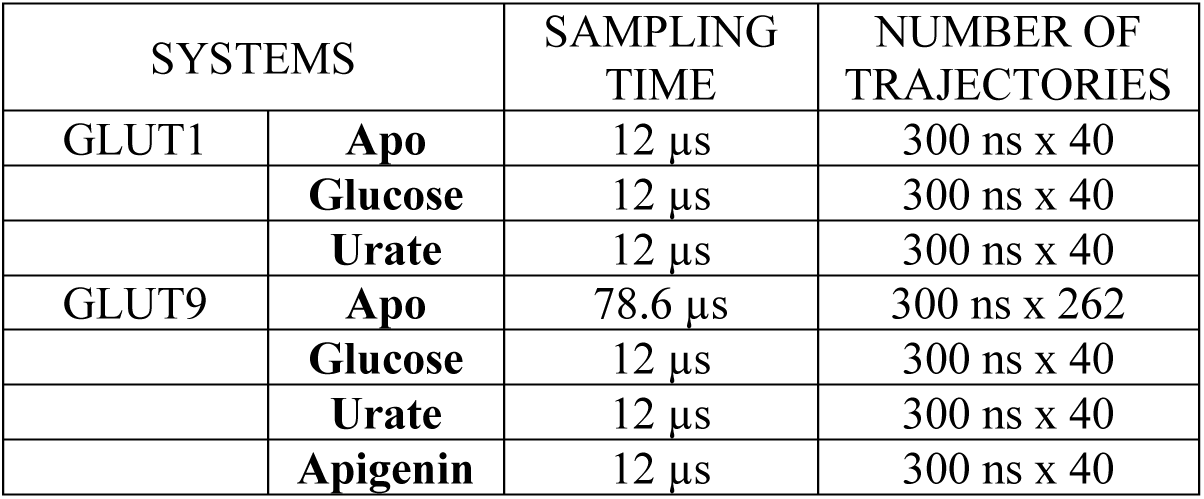
Summary of multiple adaptively sampled trajectories. Each trajectory is simulated at a time step of 0.1ns.

**Table 2.** Parameters used to build the IGME model in each system.

| System | tICA lag | Components | Clustering | MSM lag | L | $\tau_k$ |
| --- | --- | --- | --- | --- | --- | --- |
| GLUT1-Apo | 1 ns | 3 | 100 | 5 ns | 9.5 ns | 0.4 ns |
| GLUT9-Apo | 0.2 ns | 3 | 100 | 2 ns | 9.6 ns | 0.4 ns |
| GLUT9-API | 2 ns | 3 | 100 | 2 ns | 9.5 ns | 0.1 ns |

A 40ns NPT equilibration run at 310K was performed for each system, with 5 kcal/mol group restraints applied to the ligands with a box of 12×12×12 as a flat bottom to prevent the ligand from leaving the binding site during equilibration. The rest of the settings were kept the same as the default equilibration protocol in HTMD. In the production protocol, we ran the simulations in the NVT ensemble using a Langevin thermostat and damping constant of 0.1 ps^-1^. Hydrogen mass repartitioning scheme was enabled to implement a 4 fs timestep. During the production phase, all restraints were removed. For another set of simulations, the restraints box was expanded to allow the ligand to move inside the channel but not to diffuse into the solvent. An Adaptive Bandit-enhanced sampling protocol utilizing multiple short simulations based on Markov state models (MSMs) was employed. This adaptive algorithm iteratively conducts short parallel simulations, minimizing redundancy by discretizing the conformational space into an MSM and estimating free energy from the stationary distribution of each state. Simulations were restarted from low-energy conformations. The MetricSelfDistance function assessed native Cα contacts across residues to construct the MSMs, using an exploration value of 0.01 and a goal-scoring function of 0.3. Each round consisted of four 100 ns simulations, with trajectory frames recorded every 0.1 ns. The GLUT9 apo system was simulated for a cumulative 78.6 µs to achieve enough sampling to describe the opening and closure of either side of the transporter, while 12 µs was enough to separate conformations for other systems. All simulations were carried out using the ACEMD molecular dynamics engine^21, 26^.

### Non-Markov State Model

The Integrative Generalized Master Equation (IGME) based non-Markovian dynamic model was built based on a published IGME tutorial^13^. Non-Markovian models like quasi-MSM overcome the data challenge of Markovian models, which require a long enough lag time to allow transition between states to become Markovian, by using the memory kernels^27^. This allows much more precise predictions of slow dynamics from significantly short MD simulations^27^. However, numerical instability becomes the major challenge of the qMSM. The recently introduced new IGME model solves this issue through the utilization of the integrals of memory kernels, thus reducing the numerical fluctuation^28^. Two hyperparameters were used to build the IGME model, τ_*k*_ and *L*. τ_*k*_ is the memory kernel relaxation time, which should be long enough for the memory kernels to decay to zero. *L* is the length of the input data used in least-squares fitting. To fine-tune the hyperparameters, 100 IGME models were scanned for each system and the best hyperparameters were selected based on the calculated root mean squared error (RMSE) (Figure S12).

In this study, the χ_1_ angle of essential ligand binding residues and distance between helix were used as the input feature of the IGME model (details see supplementary). Featurized trajectories were then projected onto three independent components (ICs) using time-lagged independent component analysis (tICA). An appropriate tICA lag time that gives a good separation on the projection was then selected for every system. K-means was then used to further cluster the data into 100 clusters based on the selected ICs in every system. The IGME model starts with a microstate MSM using the clustered data. Then PCCA+ kinetic lumping was applied to convert the microstate MSM into a macrostate MSM with a selected number of states for every system. IGME then predicts a more accurate transition probability matrix (TPM) from the TPM of the macrostate MSM. The Chapman-Kolmogorov test was used to assess the quality of the IGME models. The mean first passage times (MFPT) between states were calculated with the row-normalized TPM. The net flux pathways between macrostates, which start from the highest energy state to the lowest energy state, were conculcated using the transition pathway theory (TPT) function. For each metastable state, a structure that is closest to the k-means clustering centre was selected and extracted as the representative conformation of this state. Moreover, all the frames belonging to each state were selected and merged using MDAnalysis for further analysis^29, 30^. Table S2 represents all the parameters used in generating the IGME model.

### PathDetect-SOM

During the MD simulation, glucose and urate exit the transporter via the intracellular side in both GLUT1 and GLUT9. Pathway analysis of these ligands was carried out using an artificial neuron network-based method called PathDetect-SOM^14^. To train the model, residues that surround the intracellular side of the transporters were selected (details in supplementary). The time-dependent pairwise Euclidean distance between the selected residues and the ligand would be automatically computed. This was then used as the input feature which was used to train the self-organizing map (SOM) model iteratively. Each frame was considered as a data point and assigned to the most familiar neuron. These neurons were then grouped into an optimal number of clusters based on the silhouette profile. The representative structure of each neuron is saved using GROMACS, and for each cluster, the tool would identify a representative neuron. All the frames belonging to each neuron were extracted and merged using MDTraj^31^. In this study, default 10 x 10 sheet-shaped SOMs without periodic boundary conditions were trained over 5000 training cycles. The distance capping was set to 12Å, where distances exceeding the threshold would be ignored and considered as the capping value. The pathway identified by this tool can be directly illustrated by constructing the neuron transition network using igraph. The heatmap which contains the contact distance between the ligand and selected input residues across the frame of each cluster was calculated via MDAnalysis^29, 30^.

### Trajectory Analysis

Quantitative analysis of any conformational changes is based on the analysis of the whole frames extracted for each state and cluster. The χ_1_ angle distribution of the specific residue and the solvent accessible surface area (SASA) of each state are calculated using MDTraj^31^. Distance distributions between either atoms or residues are computed using MDAnalysis^29, 30^. MDciao^32^, is utilized to investigate any changes in the residue-residue contact frequencies, residence times and χ_1_ angle orientations.

## Notes

### Competing Interest Statement

The authors have declared no competing interest.

## References

(1) Mueckler, M.; Thorens, B. The SLC2 (GLUT) family of membrane transporters. Mol Aspects Med 2013, 34 (2-3), 121–138. DOI: 10.1016/j.mam.2012.07.001

(2) Thorens, B.; Mueckler, M. Glucose transporters in the 21st Century. Am J Physiol Endocrinol Metab 2010, 298 (2), E141–145. DOI: 10.1152/ajpendo.00712.2009

(3) Bibert, S.; Hess, S. K.; Firsov, D.; Thorens, B.; Geering, K.; Horisberger, J. D.; Bonny, O. Mouse GLUT9: evidences for a urate uniporter. Am J Physiol Renal Physiol 2009, 297 (3), F612–619. DOI: 10.1152/ajprenal.00139.2009

(4) Dehghan, A.; Kottgen, A.; Yang, Q.; Hwang, S. J.; Kao, W. L.; Rivadeneira, F.; Boerwinkle, E.; Levy, D.; Hofman, A.; Astor, B. C.;, et al. Association of three genetic loci with uric acid concentration and risk of gout: a genome-wide association study. Lancet 2008, 372 (9654), 1953–1961. DOI: 10.1016/S0140-6736(08)61343-4

(5) Shen, Z.; Xu, L.; Wu, T.; Wang, H.; Wang, Q.; Ge, X.; Kong, F.; Huang, G.; Pan, X. Structural basis for urate recognition and apigenin inhibition of human GLUT9. Nat Commun 2024, 15 (1), 5039. DOI: 10.1038/s41467-024-49420-9

(6) Nomura, N.; Verdon, G.; Kang, H. J.; Shimamura, T.; Nomura, Y.; Sonoda, Y.; Hussien, S. A.; Qureshi, A. A.; Coincon, M.; Sato, Y.;, et al. Structure and mechanism of the mammalian fructose transporter GLUT5. Nature 2015, 526 (7573), 397–401. DOI: 10.1038/nature14909

(7) Deng, D.; Xu, C.; Sun, P.; Wu, J.; Yan, C.; Hu, M.; Yan, N. Crystal structure of the human glucose transporter GLUT1. Nature 2014, 510 (7503), 121–125. DOI: 10.1038/nature13306

(8) Anzai, N.; Ichida, K.; Jutabha, P.; Kimura, T.; Babu, E.; Jin, C. J.; Srivastava, S.; Kitamura, K.; Hisatome, I.; Endou, H.; Sakurai, H. Plasma urate level is directly regulated by a voltage-driven urate efflux transporter URATv1 (SLC2A9) in humans. J Biol Chem 2008, 283 (40), 26834–26838. DOI: 10.1074/jbc.C800156200

(9) Dror, R. O.; Dirks, R. M.; Grossman, J. P.; Xu, H.; Shaw, D. E. Biomolecular simulation: a computational microscope for molecular biology. Annu Rev Biophys 2012, 41, 429–452. DOI: 10.1146/annurev-biophys-042910-155245

(10) Hollingsworth, S. A.; Dror, R. O. Molecular Dynamics Simulation for All. Neuron 2018, 99 (6), 1129–1143. DOI: 10.1016/j.neuron.2018.08.011

(11) Chodera, J. D.; Noe, F. Markov state models of biomolecular conformational dynamics. Curr Opin Struct Biol 2014, 25, 135–144. DOI: 10.1016/j.sbi.2014.04.002

(12) Prinz, J. H.; Wu, H.; Sarich, M.; Keller, B.; Senne, M.; Held, M.; Chodera, J. D.; Schutte, C.; Noe, F. Markov models of molecular kinetics: generation and validation. J Chem Phys 2011, 134 (17), 174105. DOI: 10.1063/1.3565032

(13) Wu, Y.; Cao, S.; Qiu, Y.; Huang, X. Tutorial on how to build non-Markovian dynamic models from molecular dynamics simulations for studying protein conformational changes. J Chem Phys 2024, 160 (12). DOI: 10.1063/5.0189429

(14) Motta, S.; Callea, L.; Bonati, L.; Pandini, A. PathDetect-SOM: A Neural Network Approach for the Identification of Pathways in Ligand Binding Simulations. J Chem Theory Comput 2022, 18 (3), 1957–1968. DOI: 10.1021/acs.jctc.1c01163

(15) Fu, X.; Zhang, G.; Liu, R.; Wei, J.; Zhang-Negrerie, D.; Jian, X.; Gao, Q. Mechanistic Study of Human Glucose Transport Mediated by GLUT1. J Chem Inf Model 2016, 56 (3), 517–526. DOI: 10.1021/acs.jcim.5b00597

(16) Park, M. S. Molecular Dynamics Simulations of the Human Glucose Transporter GLUT1. PLoS One 2015, 10 (4), e0125361. DOI: 10.1371/journal.pone.0125361

(17) Ung, P. M.; Song, W.; Cheng, L.; Zhao, X.; Hu, H.; Chen, L.; Schlessinger, A. Inhibitor Discovery for the Human GLUT1 from Homology Modeling and Virtual Screening. ACS Chem Biol 2016, 11 (7), 1908–1916. DOI: 10.1021/acschembio.6b00304

(18) Hully, M.; Vuillaumier-Barrot, S.; Le Bizec, C.; Boddaert, N.; Kaminska, A.; Lascelles, K.; de Lonlay, P.; Cances, C.; des Portes, V.; Roubertie, A.;, et al. From splitting GLUT1 deficiency syndromes to overlapping phenotypes. Eur J Med Genet 2015, 58 (9), 443–454. DOI: 10.1016/j.ejmg.2015.06.007

(19) Ruiz, A.; Gautschi, I.; Schild, L.; Bonny, O. Human Mutations in SLC2A9 (Glut9) Affect Transport Capacity for Urate. Front Physiol 2018, 9, 476. DOI: 10.3389/fphys.2018.00476

(20) Li, Y.; Zhao, Z.; Luo, J.; Jiang, Y.; Li, L.; Chen, Y.; Zhang, L.; Huang, Q.; Cao, Y.; Zhou, P.;, et al. Apigenin ameliorates hyperuricemic nephropathy by inhibiting URAT1 and GLUT9 and relieving renal fibrosis via the Wnt/beta-catenin pathway. Phytomedicine 2021, 87, 153585. DOI: 10.1016/j.phymed.2021.153585

(21) Doerr, S.; Harvey, M. J.; Noe, F.; De Fabritiis, G. HTMD: High-Throughput Molecular Dynamics for Molecular Discovery. J Chem Theory Comput 2016, 12 (4), 1845–1852. DOI: 10.1021/acs.jctc.6b00049

(22) Martinez-Rosell, G.; Giorgino, T.; De Fabritiis, G. PlayMolecule ProteinPrepare: A Web Application for Protein Preparation for Molecular Dynamics Simulations. J Chem Inf Model 2017, 57 (7), 1511–1516. DOI: 10.1021/acs.jcim.7b00190

(23) Huang, J.; Rauscher, S.; Nawrocki, G.; Ran, T.; Feig, M.; de Groot, B. L.; Grubmuller, H.; MacKerell, A. D., Jr. CHARMM36m: an improved force field for folded and intrinsically disordered proteins. Nat Methods 2017, 14 (1), 71–73. DOI: 10.1038/nmeth.4067

(24) Jorgensen, W. L.; Jenson, C. Temperature dependence of TIP3P, SPC, and TIP4P water from NPT Monte Carlo simulations: Seeking temperatures of maximum density. Journal of Computational Chemistry 1998, 19 (10), 1179–1186. DOI: 10.1002/(sici)1096-987x(19980730)19:10<1179::Aid-jcc6>3.0.Co;2-j.

(25) Vanommeslaeghe, K.; MacKerell, A. D., Jr. Automation of the CHARMM General Force Field (CGenFF) I: bond perception and atom typing. J Chem Inf Model 2012, 52 (12), 3144–3154. DOI: 10.1021/ci300363c

(26) Harvey, M. J.; Giupponi, G.; Fabritiis, G. D. ACEMD: Accelerating Biomolecular Dynamics in the Microsecond Time Scale. J Chem Theory Comput 2009, 5 (6), 1632–1639. DOI: 10.1021/ct9000685

(27) Cao, S.; Montoya-Castillo, A.; Wang, W.; Markland, T. E.; Huang, X. On the advantages of exploiting memory in Markov state models for biomolecular dynamics. J Chem Phys 2020, 153 (1), 014105. DOI: 10.1063/5.0010787

(28) Cao, S.; Qiu, Y.; Kalin, M. L.; Huang, X. Integrative generalized master equation: A method to study long-timescale biomolecular dynamics via the integrals of memory kernels. J Chem Phys 2023, 159 (13). DOI: 10.1063/5.0167287

(29) Gowers, R.; Linke, M.; Barnoud, J.; Reddy, T.; Melo, M.; Seyler, S. L.; Dotson, D.; Domanski, J.; Buchoux, S.; Kenney, I. MDAnalysis: a Python package for the rapid analysis of molecular dynamics simulations; 2016.

(30) Michaud-Agrawal, N.; Denning, E. J.; Woolf, T. B.; Beckstein, O. MDAnalysis: a toolkit for the analysis of molecular dynamics simulations. J Comput Chem 2011, 32 (10), 2319–2327. DOI: 10.1002/jcc.21787

(31) McGibbon, R. T.; Beauchamp, K. A.; Harrigan, M. P.; Klein, C.; Swails, J. M.; Hernandez, C. X.; Schwantes, C. R.; Wang, L. P.; Lane, T. J.; Pande, V. S. MDTraj: A Modern Open Library for the Analysis of Molecular Dynamics Trajectories. Biophys J 2015, 109 (8), 1528–1532. DOI: 10.1016/j.bpj.2015.08.015

(32) Pérez-Hernández, G.; Hildebrand, P. W. mdciao: Accessible Analysis and Visualization of Molecular Dynamics Simulation Data. bioRxiv 2022. DOI: 10.1101/2022.07.15.500163.

