## Supplementary Information for "Mechanistic Insights into the Substrate Egress Pathways of Human Glucose Transporters GLUT1 and GLUT9"

**Supplementary Information Text**

**Residues used in nMSM**

In this study, essential residues involved in ligand binding within GLUT1 and GLUT9 were identified and selected according to previous studies[1, 2]. These critical residues were then superimposed from structural alignment to ensure consistent selection criteria across both proteins. In other words, if a key residue in GLUT1 did not have a corresponding essential residue in GLUT9, the aligned counterpart residue would be included to maintain uniformity in residue selection between the two structures. The residue selected in GLUT1 to calculate the $X_1$ angle are: $T_{30}$, $S_{80}$, $L_{137}$, $R_{153}$, $Q_{161}$, $I_{164}$, $I_{168}$, $A_{171}$, $Q_{282}$, $Q_{283}$, $I_{287}$, $N_{288}$, $F_{291}$, $N_{317}$, $R_{333}$, $F_{379}$, $E_{380}$, $W_{388}$, $W_{412}$. The residue selected in GLUT9 to calculate the $X_1$ angle are: $L_{75}$, $T_{125}$, $L_{182}$, $R_{198}$, $A_{206}$, $F_{209}$, $V_{213}$, $G_{216}$, $Y_{327}$, $Q_{328}$, $L_{332}$, $N_{333}$, $W_{336}$, $E_{364}$, $R_{380}$, $F_{426}$, $C_{427}$, $F_{435}$, $W_{459}$.

Except these residues participate in substrate transport, several residue pairs are also selected and their distance between Cαs is used to describe the helix movements in transporters. In GLUT1, residue pairs chosen are: $G_{10}$ – $I_{272}$, $V_{32}$ – $A_{331}$, $P_{58}$ – $P_{306}$, $F_{86}$ – $R_{330}$, $R_{93}$ – $R_{334}$, $L_{115}$ – $M_{364}$, $F_{119}$ – $V_{433}$, $V_{147}$ – $L_{394}$, $R_{153}$ – $P_{399}$, $G_{175}$ – $C_{429}$, $L_{185}$ – $L_{357}$ and $F_{206}$ – $F_{450}$. While in GLUT9, the chosen residue pairs are: $S_{55}$ – $V_{313}$, $H_{97}$ – $A_{346}$, $P_{103}$ – $P_{350}$, $L_{135}$ – $H_{377}$, $K_{138}$ – $R_{380}$, $A_{161}$ – $W_{410}$, $F_{164}$ – $C_{480}$, $I_{192}$ – $F_{441}$, $K_{195}$ – $P_{448}$, $G_{220}$ – $L_{476}$, $T_{230}$ – $L_{404}$ and $L_{249}$ – $V_{498}$.

### Supplementary Figures

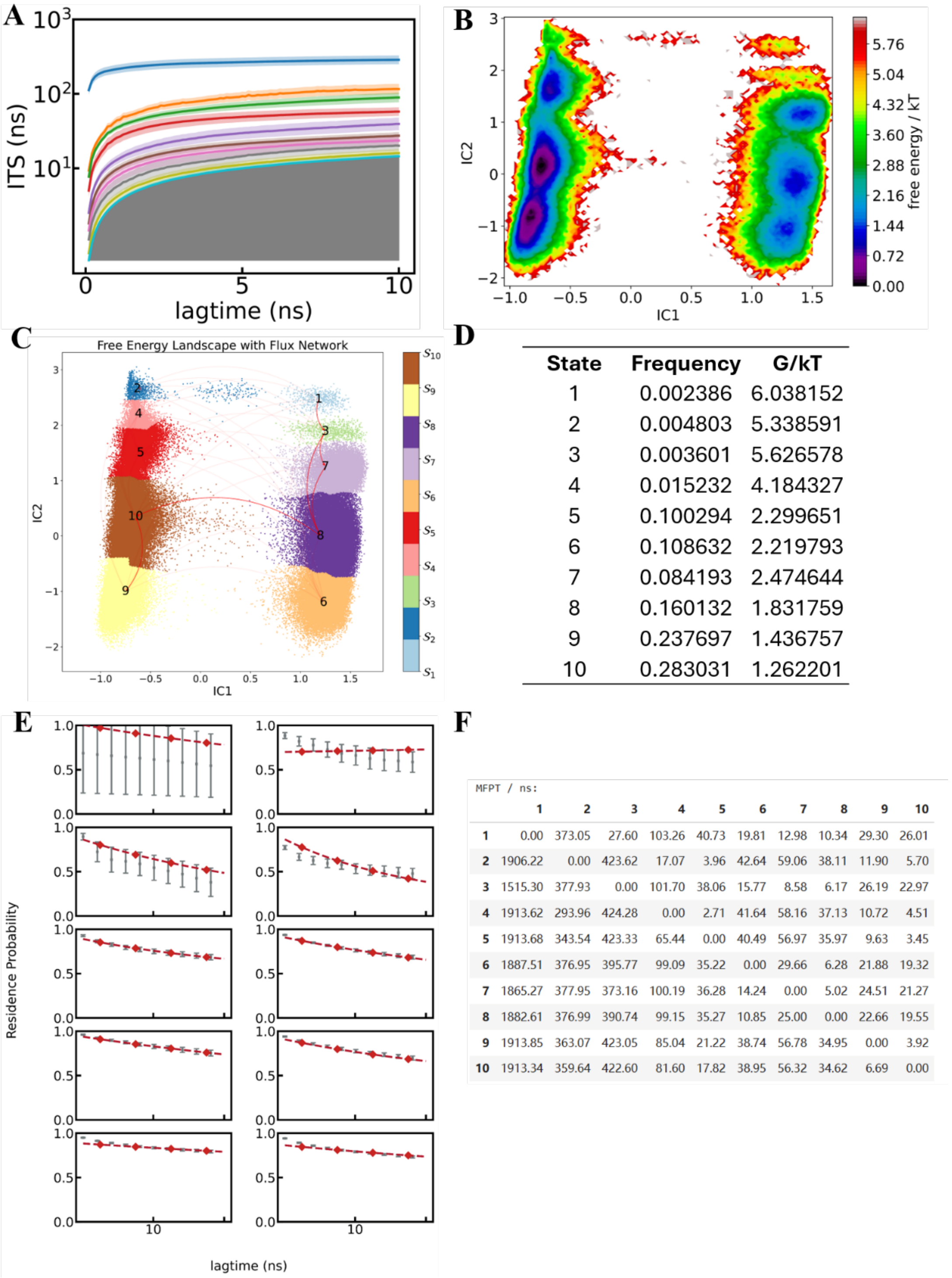


D

| State | Frequency | G/kT |
|---|---|---|
| 1 | 0.002386 | 6.038152 |
| 2 | 0.004803 | 5.338591 |
| 3 | 0.003601 | 5.626578 |
| 4 | 0.015232 | 4.184327 |
| 5 | 0.100294 | 2.299651 |
| 6 | 0.108632 | 2.219793 |
| 7 | 0.084193 | 2.474644 |
| 8 | 0.160132 | 1.831759 |
| 9 | 0.237697 | 1.436757 |
| 10 | 0.283031 | 1.262201 |

F

MFPT / ns:

| | 1 | 2 | 3 | 4 | 5 | 6 | 7 | 8 | 9 | 10 |
|---|---|---|---|---|---|---|---|---|---|---|
| 1 | 0.00 | 373.05 | 27.60 | 103.26 | 40.73 | 19.81 | 12.98 | 10.34 | 29.30 | 26.01 |
| 2 | 1906.22 | 0.00 | 423.62 | 17.07 | 3.96 | 42.64 | 59.06 | 38.11 | 11.90 | 5.70 |
| 3 | 1515.30 | 377.93 | 0.00 | 101.70 | 38.06 | 15.77 | 8.58 | 6.17 | 26.19 | 22.97 |
| 4 | 1913.62 | 293.96 | 424.28 | 0.00 | 2.71 | 41.64 | 58.16 | 37.13 | 10.72 | 4.51 |
| 5 | 1913.68 | 343.54 | 423.33 | 65.44 | 0.00 | 40.49 | 56.97 | 35.97 | 9.63 | 3.45 |
| 6 | 1887.51 | 376.95 | 395.77 | 99.09 | 35.22 | 0.00 | 29.66 | 6.28 | 21.88 | 19.32 |
| 7 | 1865.27 | 377.95 | 373.16 | 100.19 | 36.28 | 14.24 | 0.00 | 5.02 | 24.51 | 21.27 |
| 8 | 1882.61 | 376.99 | 390.74 | 99.15 | 35.27 | 10.85 | 25.00 | 0.00 | 22.66 | 19.55 |
| 9 | 1913.85 | 363.07 | 423.05 | 85.04 | 21.22 | 38.74 | 56.78 | 34.95 | 0.00 | 3.92 |
| 10 | 1913.34 | 359.64 | 422.60 | 81.60 | 17.82 | 38.95 | 56.32 | 34.62 | 6.69 | 0.00 |

**Figure S1 – nMSM model of GLUT1 Apo**. **A.** Implied timescales (ITS) plot, a lag time of 5ns was chosen to build the MSM. **B.** Free energy landscape projected by independent components. **C.** State distribution with flux network. **D.** Frequency and calculated free energy of each state. **E.** Chapman-Kolmogorov (CK) test plot. **F.** Mean first passage times (MFPT) between metastable states.

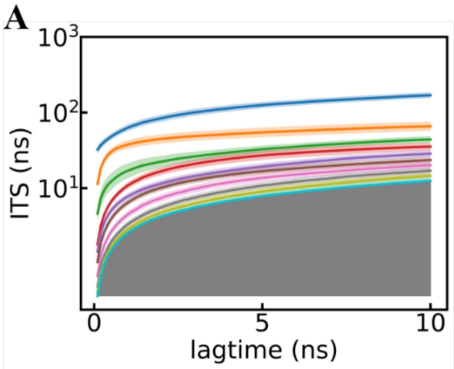


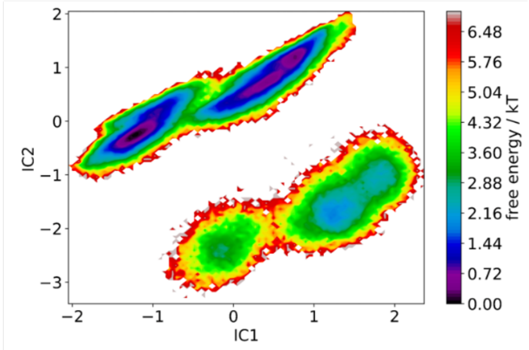


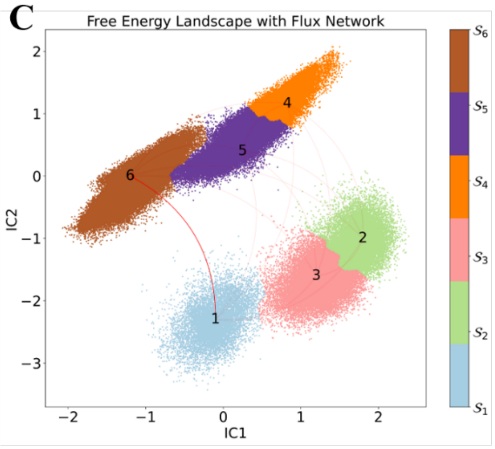


**D**

| State | Frequency | G/kT |
|---|---|---|
| 1 | 0.037004 | 3.296725 |
| 2 | 0.048813 | 3.019769 |
| 3 | 0.107617 | 2.229180 |
| 4 | 0.189058 | 1.665700 |
| 5 | 0.253163 | 1.373724 |
| 6 | 0.364346 | 1.009652 |

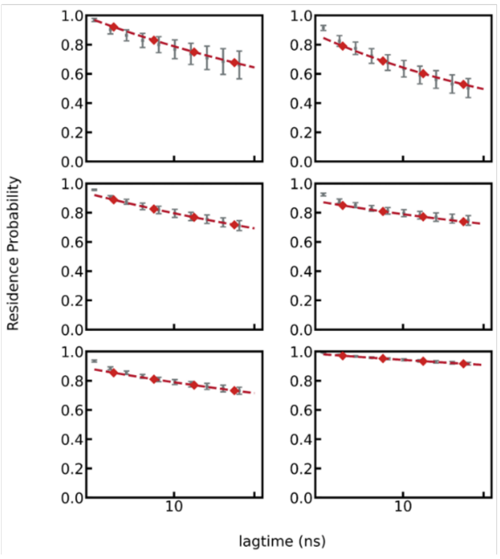


**F**

MFPT / ns:

| | 1 | 2 | 3 | 4 | 5 | 6 |
|---|---|---|---|---|---|---|
| **1** | 0.00 | 32.35 | 17.94 | 16.27 | 10.13 | 7.86 |
| **2** | 54.19 | 0.00 | 11.74 | 6.28 | 5.49 | 16.54 |
| **3** | 49.40 | 21.36 | 0.00 | 10.41 | 5.69 | 15.50 |
| **4** | 56.41 | 24.58 | 19.09 | 0.00 | 3.57 | 15.62 |
| **5** | 55.05 | 28.58 | 19.16 | 8.35 | 0.00 | 13.41 |
| **6** | 49.24 | 36.09 | 25.43 | 16.86 | 9.87 | 0.00 |

**Figure S2 – nMSM model of GLUT9 Apo. A.** Implied timescales (ITS) plot, a lag time of 5ns was chosen to build the MSM. **B.** Free energy landscape projected by independent components. **C.** State distribution with flux network. **D.** Frequency and calculated free energy of each state. **E.** Chapman-Kolmogorov (CK) test plot. **F.** Mean first passage times (MFPT) between metastable states.

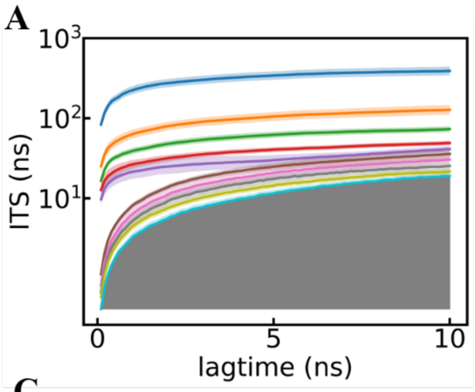


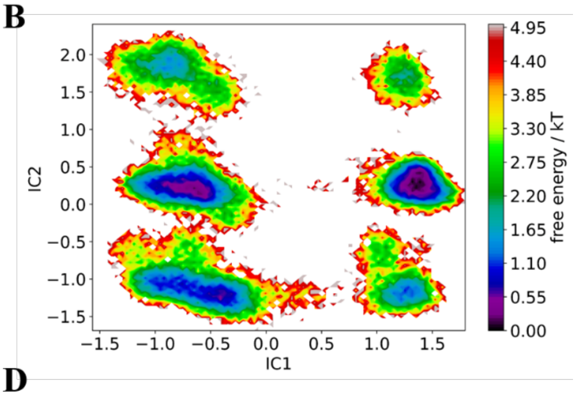


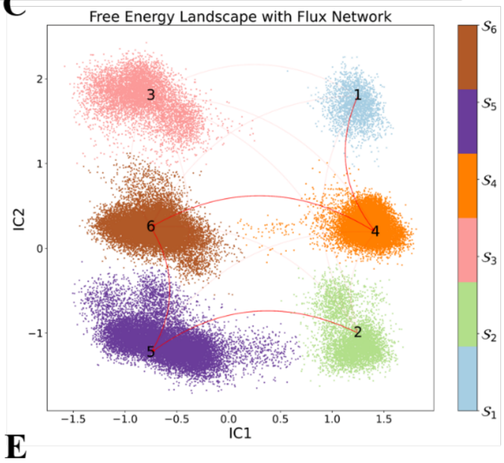


D

| State | Frequency | G/kT |
|---|---|---|
| 1 | 0.036633 | 3.306797 |
| 2 | 0.086933 | 2.442614 |
| 3 | 0.108733 | 2.218857 |
| 4 | 0.225167 | 1.490914 |
| 5 | 0.263533 | 1.333575 |
| 6 | 0.279000 | 1.276543 |

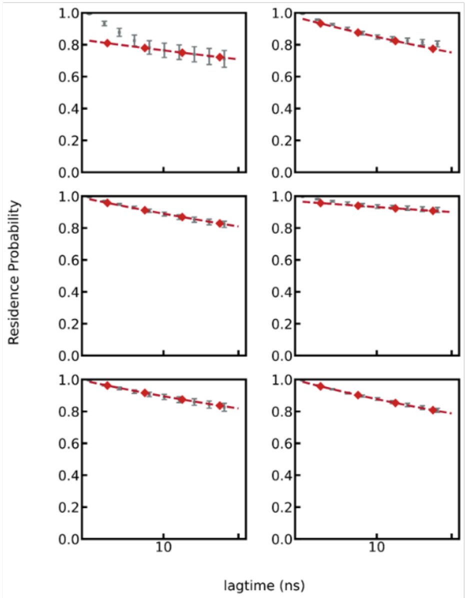


F

MFPT / ns:

| | 1 | 2 | 3 | 4 | 5 | 6 |
|---|---|---|---|---|---|---|
| **1** | 0.00 | 6.08 | 13.59 | 1.15 | 6.47 | 6.76 |
| **2** | 17.00 | 0.00 | 11.73 | 7.41 | 3.52 | 4.86 |
| **3** | 23.14 | 10.37 | 0.00 | 13.16 | 3.22 | 1.14 |
| **4** | 10.88 | 6.22 | 13.33 | 0.00 | 6.30 | 6.49 |
| **5** | 21.97 | 8.11 | 9.16 | 12.08 | 0.00 | 2.28 |
| **6** | 22.38 | 9.57 | 7.21 | 12.38 | 2.40 | 0.00 |

**Figure S3 – nMSM model of GLUT9 API. A.** Implied timescales (ITS) plot, a lag time of 5ns was chosen to build the MSM. **B.** Free energy landscape projected by independent components. **C.** State distribution with flux network. **D.** Frequency and calculated free energy of each state. **E.** Chapman-Kolmogorov (CK) test plot. **F.** Mean first passage times (MFPT) between metastable states.

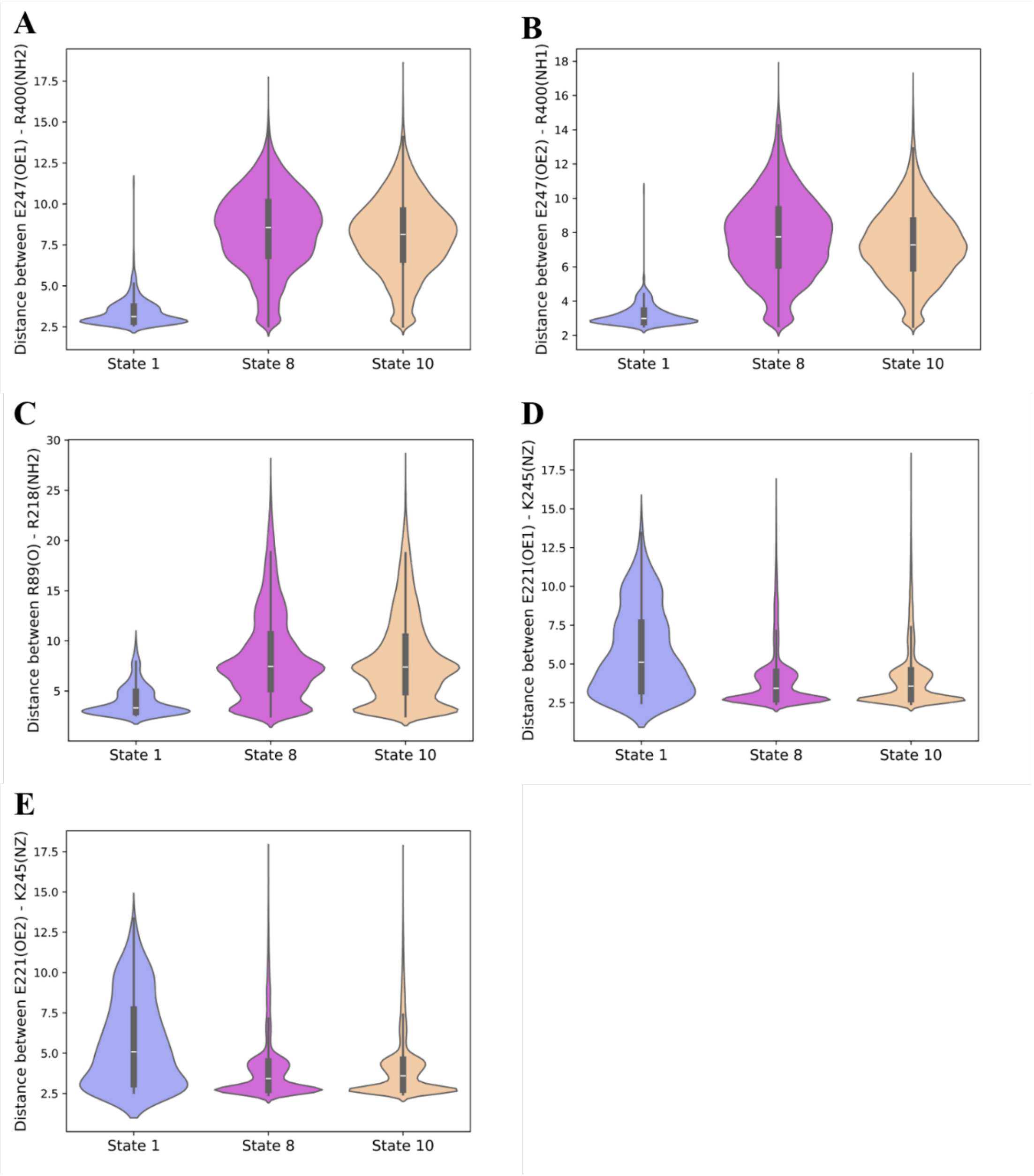


**Figure S4 – Violin plot of distance distribution between atoms of state 1, state 8, and state 10 (GLUT1 Apo). A.** Distance between $E_{247}$(OE1) and $R_{400}$(NH2). **B.** Distance between $E_{247}$(OE2) and $R_{400}$(NH1). **C.** Distance between $R_{89}$(O) and $R_{218}$(NH2). **D.** Distance between $E_{221}$(OE1) and $K_{245}$(NZ). **E.** Distance between $E_{221}$(OE2) and $K_{245}$(NZ).

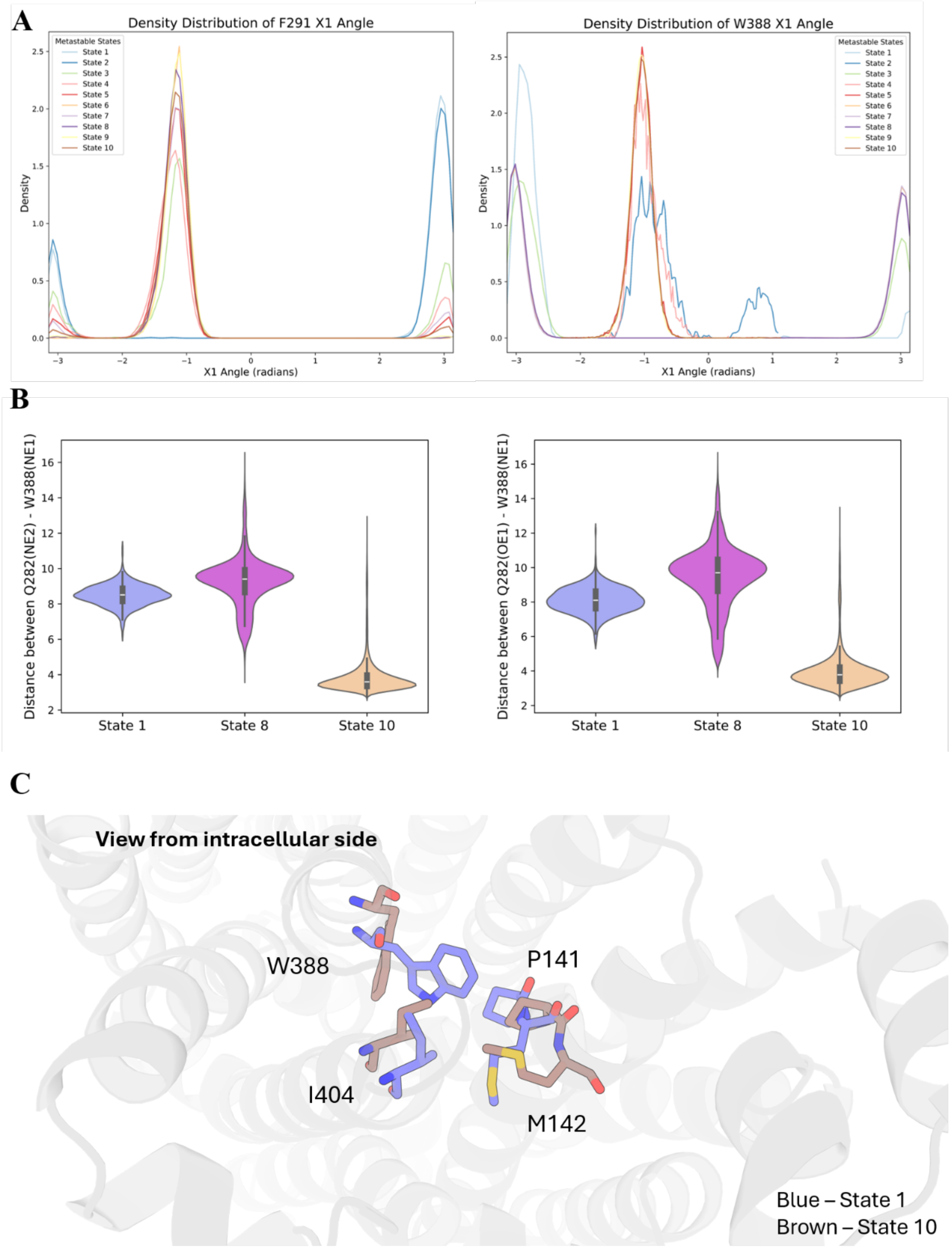


**Figure S5 – Additional figures for GLUT1 Apo. A.** The distribution of the $X_1$ angle of $F_{291}$. **B.** The distribution of the $X_1$ angle of $W_{388}$. **C.** Distance distribution of $Q_{282}$(NE2) and $Q_{282}$(OE2) to $W_{388}$(NE1). State 10 shows strong interactions between these two residues. **D.** The hydrophobic gate formed by $W_{388}$, $P_{141}$, $M_{142}$, and $I_{404}$. Blue represents the conformation in state 1 and brown represents the conformation in state 10. State 10 is showing a wider tunnel with the sidechain of $W_{388}$ being lifted towards the extracellular side.

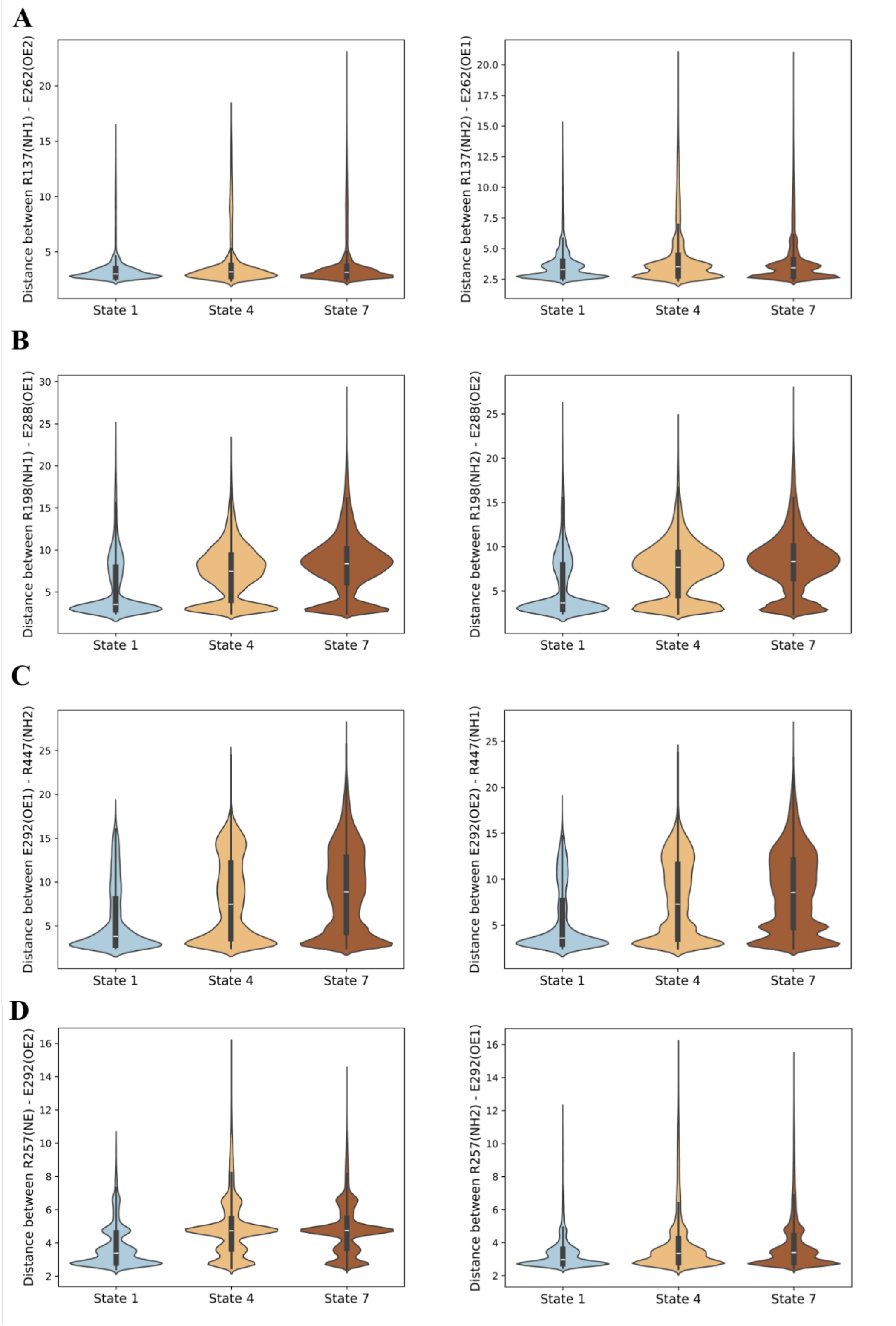


**Figure S6 – Distance distribution between residues in each state (GLUT9 Apo). A.** $R_{137}$ and $E_{262}$. **B.** $R_{198}$ and $E_{288}$. **C.** $E_{292}$ and $R_{447}$. **D.** $R_{257}$ and $E_{292}$.

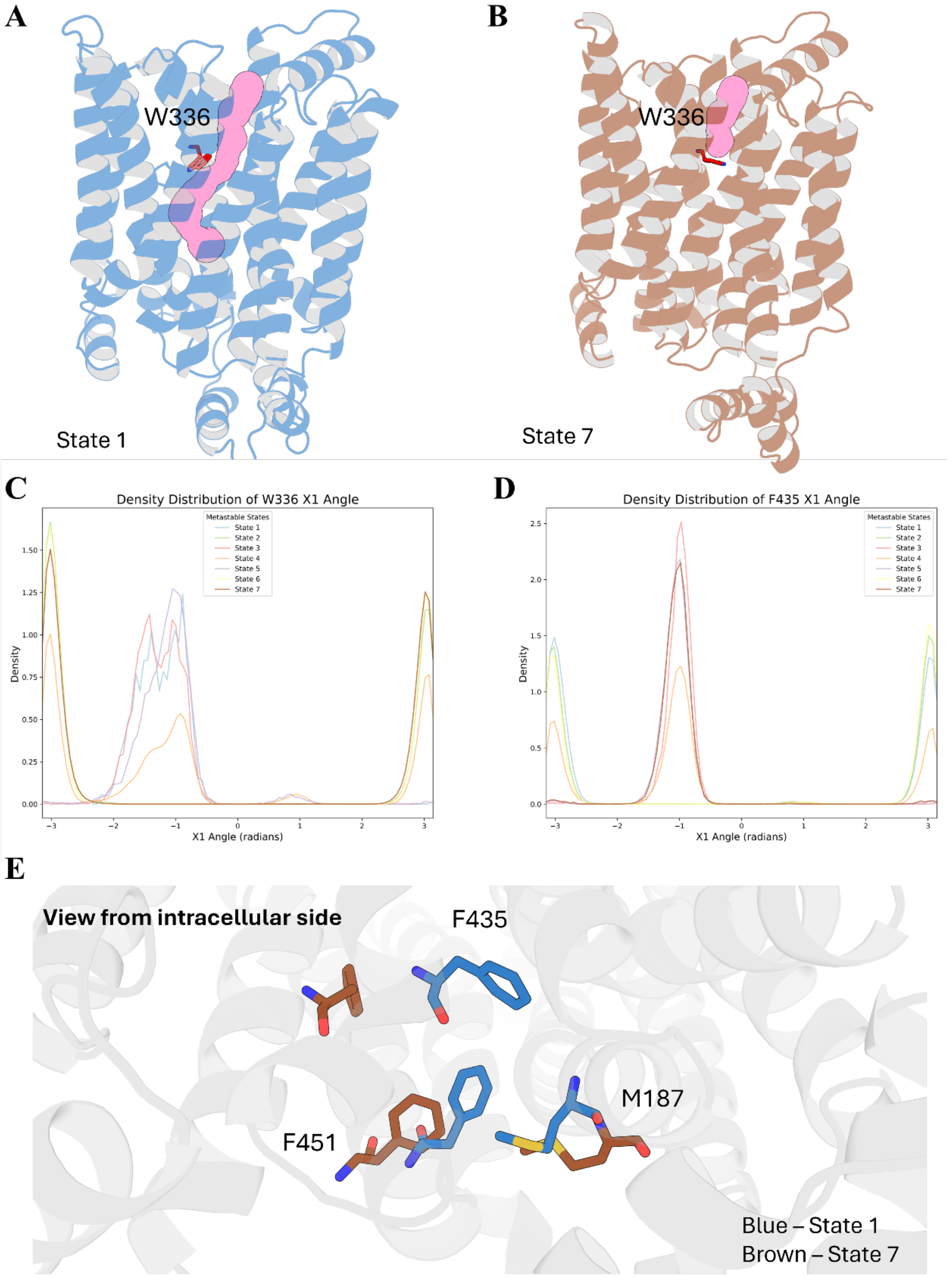


**Figure S7 – Additional figures for GLUT9 Apo. A&B.** Tunnel Detected by CAVER[3] of State 1 (Blue) and State 7 (Brown). **C.** The distribution of the $X_1$ angle of $W_{336}$. **D.** The distribution of the $X_1$ angle of $F_{435}$. **E.** The hydrophobic core formed by $M_{187}$, $F_{451}$, and $F_{435}$. Blue represents the conformation in state 1 and brown represents the conformation in state 7. State 7 shows a much wider tunnel with the orientation and movement of $F_{435}$.

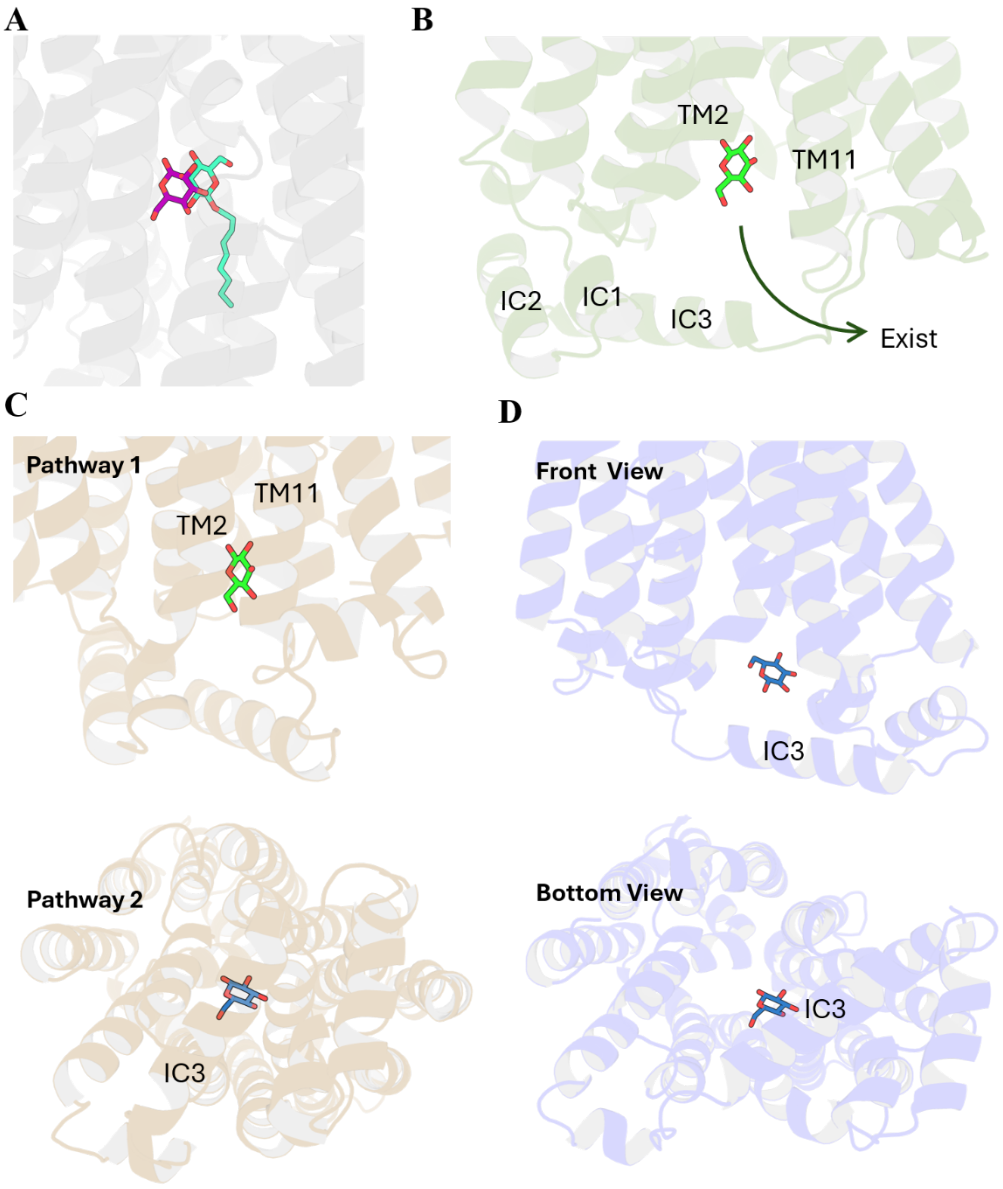


**Figure S8 – Additional figures for glucose pathway in GLUT1. A.** Glucose binding conformation comparison between the binding state (Cluster E) to the X-ray crystallography binding structure (PDB id: 4PYP). **B.** Detailed position of glucose before existence in pathway 1 (Cluster B). **C.** Superimposition of glucose to the inward close conformation determined by nMSM (state 1) to determine whether the pathway still exists when the intracellular side is closed. **D.** Front view and bottom view of the position of glucose before existence in pathway 2 (Cluster G).

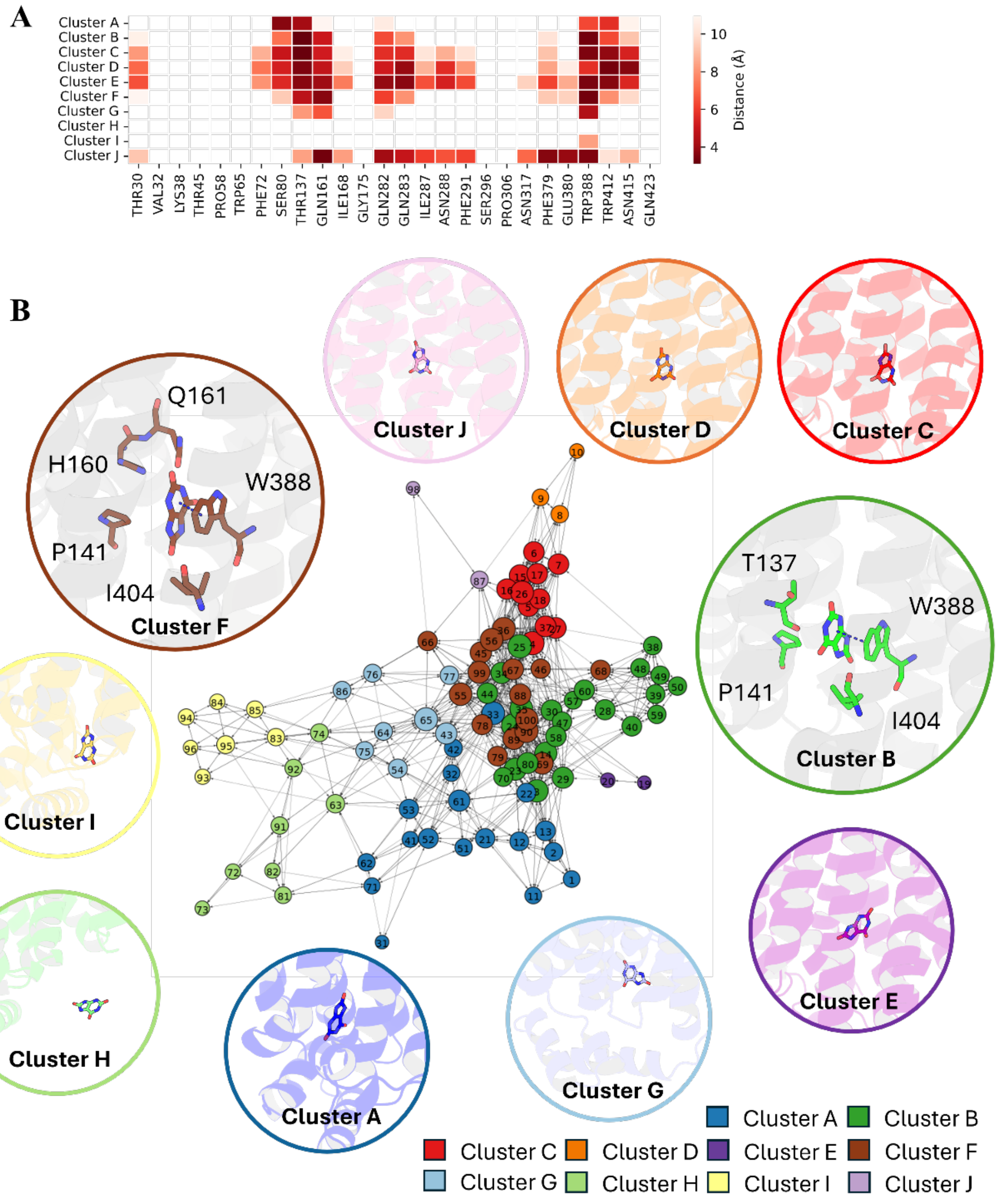


**Figure S9 – GLUT1 urate pathway. A.** Distance heatmap between urate and selected input residues during SOM training. B. Neuron transition network. Detailed ligand-protein contacts in each cluster are placed close to the neuron.

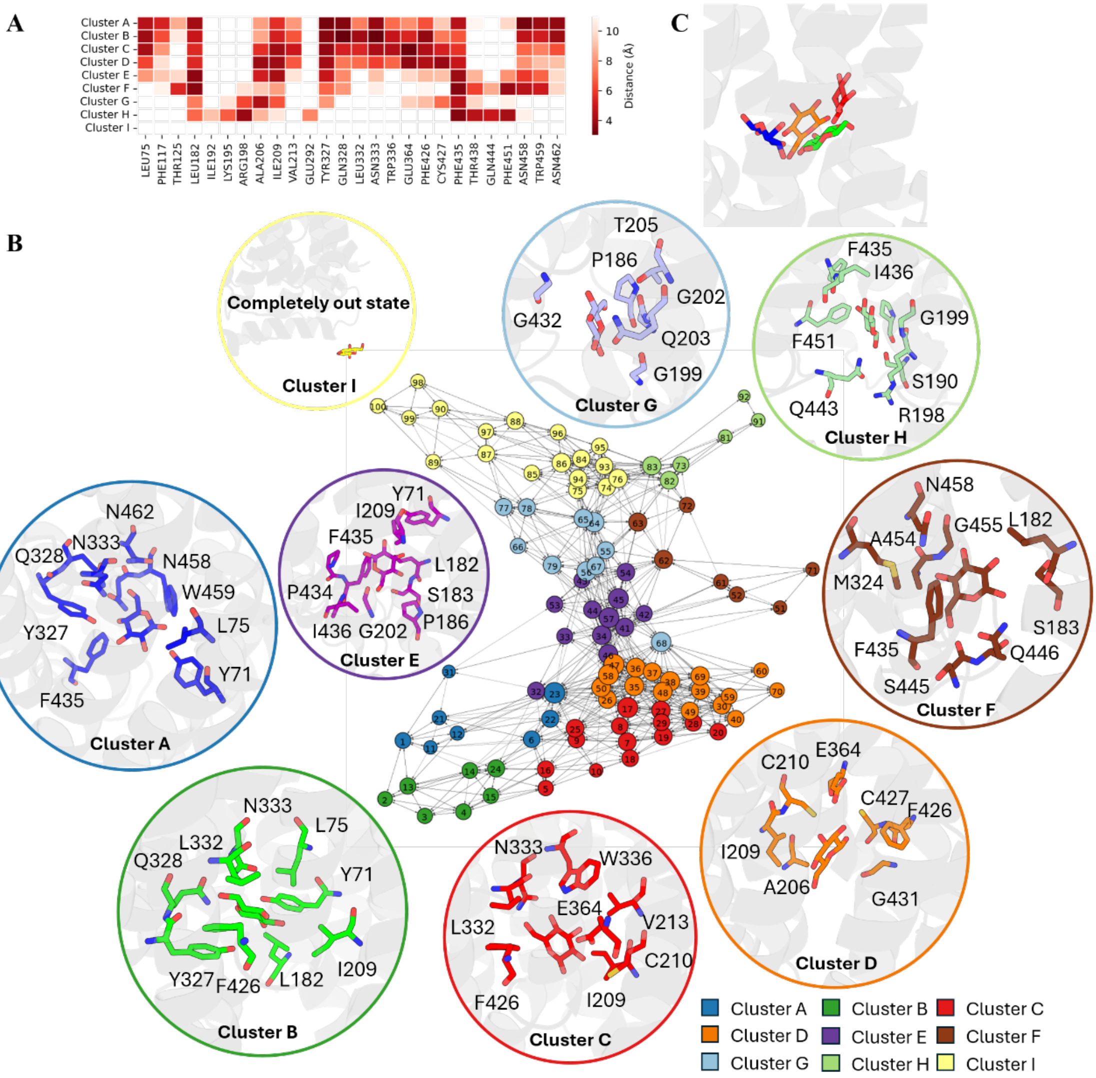

**Figure S10 – GLUT9 glucose pathway. A.** Distance heatmap between glucose and selected input residues during SOM training. **B.** Neuron transition network. Detailed ligand-protein contacts in each cluster are placed close to the neuron. **C.** Superimposition of glucose position in clusters A, B, C, and D.

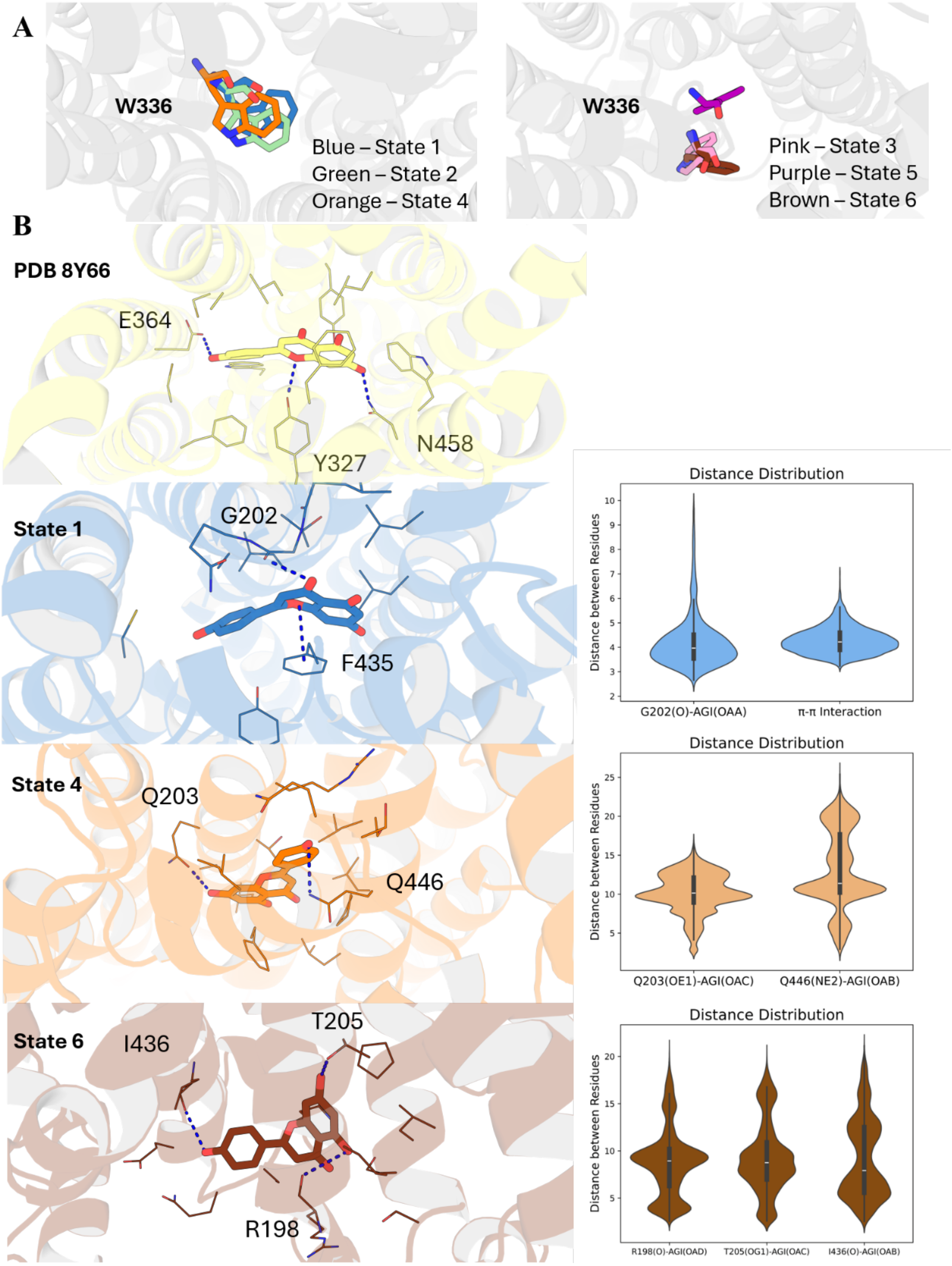


**Figure S11 – GLUT9 API MSM additional figures. A.** Conformation of W336 in each state. **B.** Ligand binding position of Apigenin in the extracted representative structure of state 1, state 4, and state 6 with the calculated distance distribution among all the frames of each state.

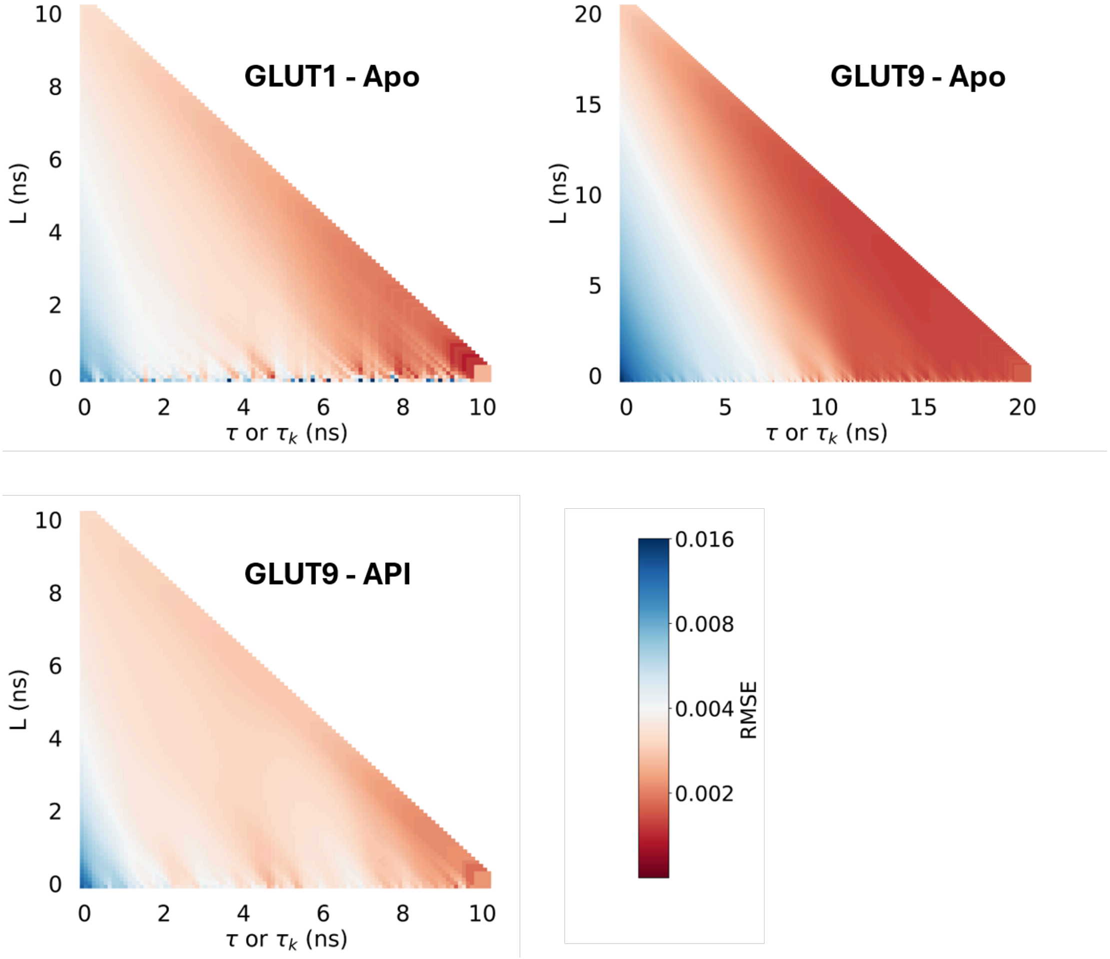


**Figure S12** – Calculated root mean squared error (RMSE) of each IGME model.

**Reference**

(1) Deng, D.; Xu, C.; Sun, P.; Wu, J.; Yan, C.; Hu, M.; Yan, N. Crystal structure of the human glucose transporter GLUT1. *Nature* **2014**, *510* (7503), 121-125. DOI: 10.1038/nature13306

(2) Shen, Z.; Xu, L.; Wu, T.; Wang, H.; Wang, Q.; Ge, X.; Kong, F.; Huang, G.; Pan, X. Structural basis for urate recognition and apigenin inhibition of human GLUT9. *Nat Commun* **2024**, *15* (1), 5039. DOI: 10.1038/s41467-024-49420-9

(3) Chovancova, E.; Pavelka, A.; Benes, P.; Strnad, O.; Brezovsky, J.; Kozlikova, B.; Gora, A.; Sustr, V.; Klvana, M.; Medek, P.; et al. CAVER 3.0: a tool for the analysis of transport pathways in dynamic protein structures. *PLoS Comput Biol* **2012**, *8* (10), e1002708. DOI: 10.1371/journal.pcbi.1002708
